# Genetic dysregulation of gene expression and splicing during a ten-year period of human aging

**DOI:** 10.1101/519520

**Authors:** Brunilda Balliu, Matthew Durrant, Olivia de Goede, Nathan Abell, Xin Li, Boxiang Liu, Michael J Gloudemans, Naomi L. Cook, Kevin S. Smith, Mauro Pala, Francesco Cucca, David Schlessinger, Siddhartha Jaiswal, Chiara Sabatti, Lars Lind, Erik Ingelsson, Stephen B Montgomery

## Abstract

Molecular and cellular changes are intrinsic to aging and age-related diseases. Prior cross-sectional studies have investigated the combined effects of age and genetics on gene expression and alternative splicing; however, there has been no long-term, longitudinal characterization of these molecular changes, especially in older age. We performed RNA sequencing in whole-blood from the same individuals from the PIVUS study at ages 70 and 80 to quantify how gene expression, alternative splicing, and their genetic regulation are altered during this 10-year period of advanced aging. We observe that individuals are more similar to their own expression profiles later in life than profiles of other individuals their own age; 93% of samples cluster with their own measurement at another age, and there is a strong correlation of genetic effects on expression between the two ages (median *ρ*_*G*_ = 0.96). Despite this, we identify 1,291 and 294 genes differentially expressed and alternatively spliced with age, as well as 529 genes with outlying individual trajectories of aging. Further, 7.8% and 9.6% of tested genes show a reduction in genetic associations with expression and alternative splicing in older age, with impacted genes enriched in DNA repair pathways. Together these findings indicate that, although gene expression and alternative splicing and their genetic regulation are mostly stable late in life, a small subset of genes is dynamic and is characterized by changes in expression and splicing and a reduction in genetic regulation.

## 1. Introduction

While an individual’s genome sequence is mostly stable throughout life, gene expression and genetic regulation of expression fluctuate in response to different environmental exposures**^? ? ?^**. The impact of aging on gene expression and genetic regulation has been well-studied in model systems, such as yeast, fruit fly or worm**^? ?^** while much less is known about the transient nature of gene regulation and expression in humans. The majority of studies that have been performed in humans have been cross-sectional**^? ? ? ? ? ? ? ? ? ?^**. The few longitudinal studies have either focused on a specific disease or intervention**^? ? ? ? ? ?^**, looked over a short time span**^?^**, or focused on cell lines**^?^**. Even less is known about the effect of age on alternative splicing and its genetics, even though changes in alternative splicing have previously been linked to aging-associated phenotypes**^? ? ? ? ?^**. As a result, a complete picture of the long-term effect of aging on gene expression and splicing and their genetic regulation in humans is still lacking.

Here we present the first long-term, longitudinal characterization of changes in gene expression and alternative splicing and their genetic regulation as a function of aging late in life. We performed RNA sequencing in whole-blood from 65 healthy participants from the PIVUS study**^?^** at both age 70 and 80, a period of the aging process characterized by high morbidity and mortality. We quantified how total and allele-specific gene expression, alternative splicing, and genetic regulation (expression and splicing quantitative trait loci; eQTLs, sQTLs) are altered over this 10-year period.

We observe that individuals are more similar to their own gene expression profiles than profiles of other individuals their own age; 93% of samples cluster with their own measurements at another age. Despite this, we identified hundreds of genes with differential expression and alternative splicing with age, as well as outlying individual trajectories of aging, i.e. individuals with extreme increase or decrease in expression with age. Moreover, we observed a strong correlation of genetic effects on expression between the two ages (*ρ*_*G*_ = .96; median across genes) and that 7.8% of genes were characterized by genetic dysregulation over the two time points. In contrast, overall allelic imbalance within an individual increases with age by 2.69% (median across individuals). Together these findings indicate that, although gene expression and alternative splicing and their genetic regulation are mostly stable late in life, a small subset of genes is dynamic and is characterized by changes in expression and splicing and a reduction in genetic regulation. The strong correlation of genetic effects and the increase in allelic imbalance with age suggests that increasing environmental variance as opposed to decreased genetic variance underlies, in part, the reduction in genetic regulation.

## 2. Results

### 2.1. Population-level age-specific expression across the transcriptome

In order to quantify the stability of gene expression levels within individuals, we measured the correlation of expression across genes between the two timepoints, after correcting for major components of gene expression variability, unrelated to age, such as technical sequencing factors and methylation-based estimates of cell type composition. We identified a moderate correlation of gene expression within an individual across genes (Spearman’s *ρ* = .30; median across individuals; Fig. S4A) and a high similarity of expression profiles; measurements of the same individual at the two ages cluster together for the 93% of samples (Fig. 1A and S4 B).

**Figure 1:**
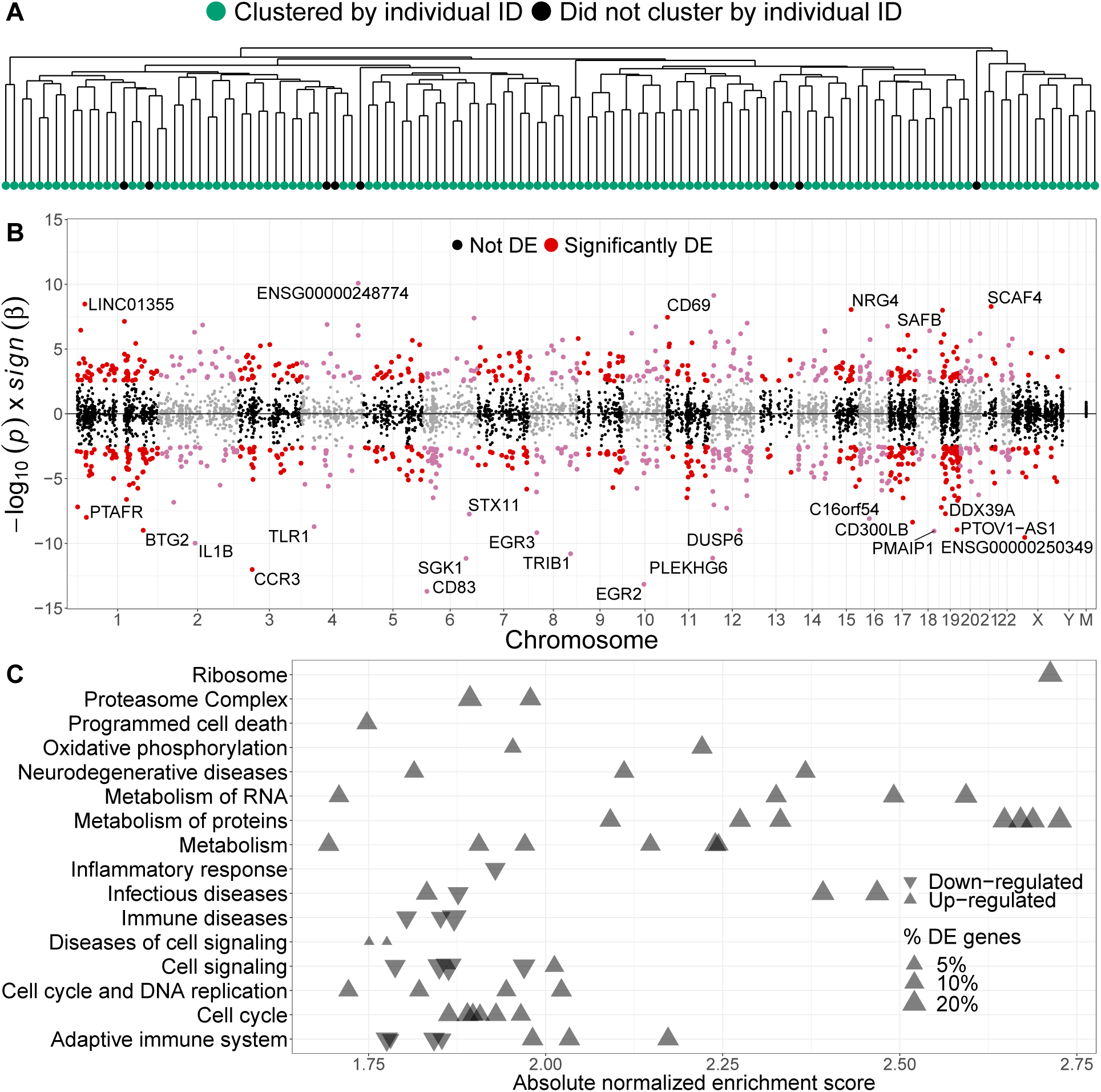
Population-level age-specific expression across the transcriptome. **(A)** Dendrogram of expression-based sample-to-sample distance. Measurements of the same individual at the two ages cluster together (green) for 93% of samples. **(B)** Mirror Manhattan plot of the expression-age discoveries. We find 1,291 age-associated genes (8% of tested genes; *FDR* ≤ 5%). Each dot represents a gene; the x-axis gives the position of the gene in the genome and the y-axis represents the direction of the age effect and the strength of the association. **(C)** Gene-set enrichment analysis of genes with age-specific expression. We observe strong enrichment for multiple age-related pathways. Each point represents a pathway; the x-axis gives the absolute enrichment score, which reflects the degree to which each pathway is over-represented at the top or bottom of the ranked list of differentially expressed genes, normalized to account for differences in gene set size and in correlations between gene sets and the expression data set. The y-axis lists the parent node of the most enriched pathways (*FDR* ≤ 5%). The names of each significant pathway are listed in Sup Files.

We investigated transcriptome-wide changes in gene expression with age (Fig. 1B and S5A and Tab. S2) and discovered 1,291 genes (8% of tested genes) whose expression levels were significantly associated with age (FDR ≤ 5%), with a slightly larger proportion of differentially expressed (DE) genes showing down-regulation with age (*π*_*down*_ = 54.29%, *χ*^2^-test; p-value = 2.2 × 10^−3^). The DE genes showed significant enrichment (FDR≤ 5%) for multiple aging-related pathways (Fig. 1C and Tab. S3), e.g. metabolism of proteins**^? ?^**, oxidative phosphorylation**^?^**, and DNA replication**^?^**. Moreover, 18 of these DE genes are previously known to be complex-trait associated genes where gene expression levels modulate disease risk in whole blood (Sup Methods and Tab. S4).

To further quantify the relative effect of age on gene expression, we estimated the proportion of expression variance (removing background noise) explained by age (Fig. S5B). Age explained 1.5% of expression variance for genes significantly associated with age. This estimate is smaller than the estimate from the uncorrected analysis of our data (average across DE genes = 7.9%) but comparable to estimates from other aging studies in humans, e.g. 2.2% in **?**.

We validated our DE genes *in silico* using summary statistics from two large cross-sectional studies of aging in human PBMCs (CHARGE**^?^**) and whole blood (SardiNIA**^?^**). We found a significant overlap between our top 1,000 DE genes and the top 1,000 DE genes from these other two studies (Hyper-geometric exact test; p-value=1.3 × 10^−16^; Fig. S5C). The 49 DE genes that are shared between the three studies (Tab. S5) are enriched in gene ontology (GO) terms related to adaptive immune response pathways that have previously been implicated in aging**^?^**, e.g. leukocyte cell-to-cell adhesion and terms reflecting the underlying age-related T cell biology (Fig. S5D). In addition, 22 and 17 known aging-and longevity-related genes from The Human Ageing Genomic Resources**^?^** GenAge (307 genes) and LongevityMap (212 genes) databases were included in our list of DE genes (Tab. S6).

To study the impact of cell-type composition in our differential expression results, we contrasted our list of DE genes to a list of 547 genes that distinguish 22 human hematopoietic cell phenotypes, including seven T cell types, naïve and memory B cells, plasma cells, NK cells, and myeloid subsets**^?^**. We found a minimal impact of the cell-type composition in our results; only 4.5% of all DE genes and 5% of top 100 DE genes are signature genes with cell-type specific expression.

### 2.2. Individual-level age-specific expression across the transcriptome

The longitudinal design of our study enabled us to also investigate changes in individual-level expression profiles with age. We searched for individuals with outlying age trajectories (schematically illustrated in Fig. 2A) and found 555 individual-gene outlier pairs from 529 unique genes (Fig. 2B and Tab. S7); 60% of which showed outlying decrease in expression with age, as opposed to increase, and 6% of which are also DE with age. Genes in which individuals showed an outlying decrease of expression with age showed significant enrichment (FDR ≤ 5%) for known age-related GO terms (Fig. 2C), i.e. activation of immune response**^?^** and regulation of proteolysis and peptidase activity**^?^**. In contrast, genes in which individuals showed an outlying increase of expression with age were not enriched for any specific functions. Only 5% of the age-trajectory outlier genes are signature genes with cell-type specific expression, indicating a minimal impact of the cell-type composition in our results.

**Figure 2:**
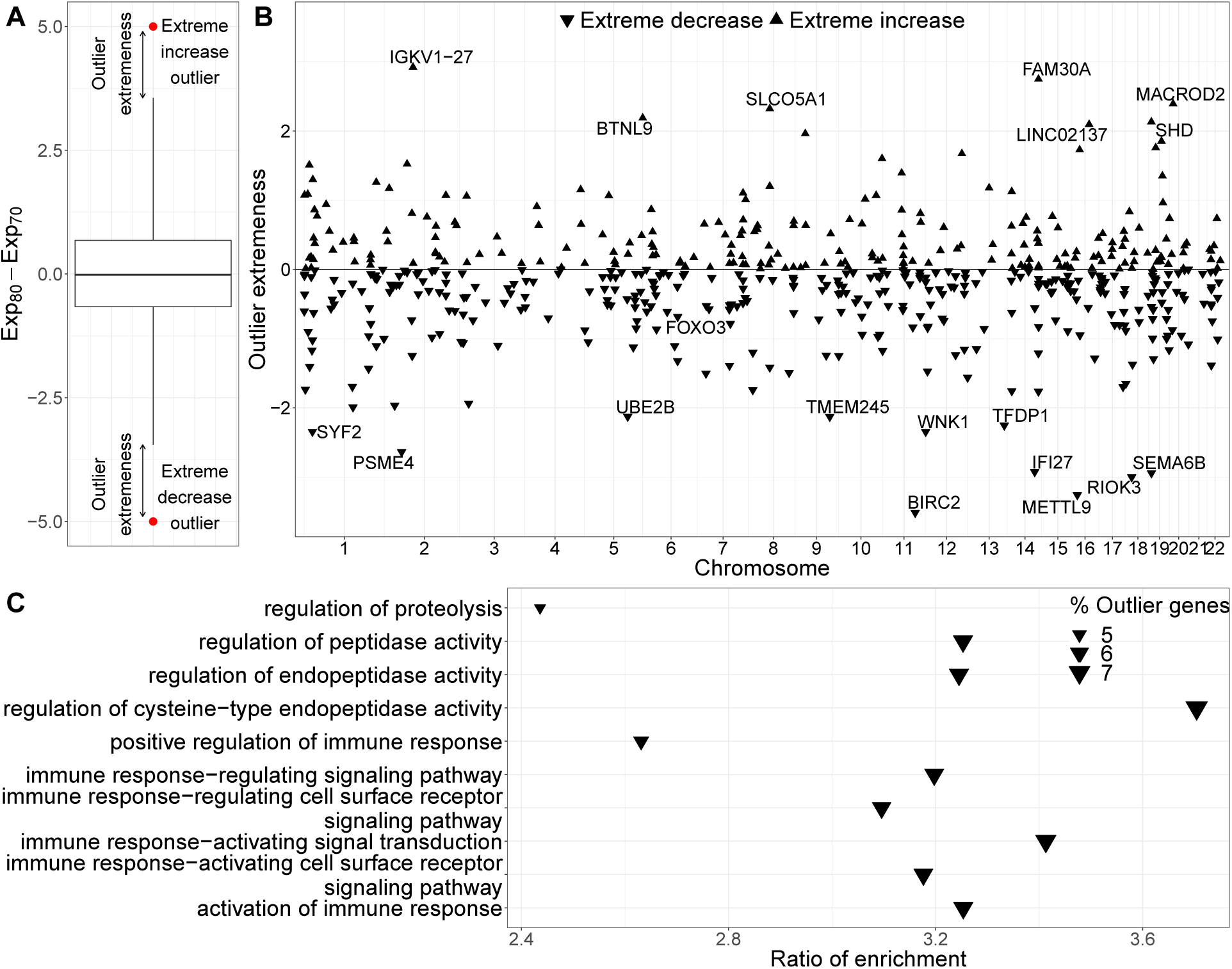
Individual-level age-specific expression across the transcriptome. **(A)** Illustration of age-trajectory outliers. Individuals are outliers for a gene if their change in the expression of the gene between the two ages falls outside the (*Q*_1_ − 3 × *IQR*, *Q*_3_ + 3 × *IQR*) range, where *Q*_1_ and *Q*_3_ are the 25th and 75th percentiles and IQR is the interquartile range. **(B)** Mirror Manhattan plot of the age-trajectory outliers. We find 555 individual-gene outlier pairs, consisting of 529 individual genes, 60% of which show extreme expression decrease with age. *IGKV1-27*, responsible for antigen binding and involved in adaptive immune response, shows the largest expression increase. *BIRC2*, a gene that inhibits apoptosis by binding to tumor necrosis factor receptor-associated factors TRAF1 and TRAF2, shows the largest expression decrease. Each dot represents a gene; the x-axis gives the position of the gene in the genome and the y-axis represents the outlier extremeness (illustrated in A). **(C)** Enrichment for GO biological processes for outlier genes. Genes with extreme expression increase with age were not enriched for any specific functions. However, genes with extreme decrease showed significant enrichment (FDR ≤ 5%) for known age-related GO terms, i.e. activation of immune response**^?^** and regulation of proteolysis and peptidase activity**^?^**.

The largest outlying expression increase with age was observed for an individual in *IGKV1-27*, a gene responsible for antigen binding and involved in adaptive immune response. The same individual was an outlier for several other immunoglobulin-related genes (Fig. S6A); we did not observe any significant changes in any of the clinical phenotypes measured at age 70 and 80 for this individual. *BIRC2*, a gene which regulates apoptosis and modulates inflammatory signaling and immunity, mitogenic kinase signaling, and cell proliferation, showed the largest expression decrease with age. The same individual was an outlier for several other genes related to proteasomal protein catabolic process, had a substantial increase in albumin levels, and was diagnosed with diabetes between age 70 and 80 (Fig. S6B, D, E). Two other individuals, one of which showed the largest increase in leukocyte counts between the two ages, were outliers for a set of genes related to adaptive immune response (Fig. S6C-E).

### 2.3. Age-specific genetic regulation of gene expression across the transcriptome

We evaluated the association between gene expression at each age group and genetic variants within 1Mb of the transcription start site using linear regression (Methods). After background noise correction (Fig. 3A and S7), we detected significant eQTLs for 1,326 genes at age 70 (8.5% of tested genes, FDR ≤ 1%, Tab. S8), 92.2% of which replicated at age 80 (FDR ≤ 10%, Fig. 3B). On the other hand, we detected eQTLs for 1,264 genes in the 80-year-old samples, 94.9% of which replicated at age 70. The depletion of genes with at least one significant eQTL (eGenes) at age 80, relative to age 70, was statistically significant (Exact McNemar’s test; p-value = 5.8 × 10^−3^). Moreover, the proportion of eGenes discovered at age 70 that replicated at age 80 was significantly smaller than the proportion of eGenes discovered at age 80 that replicated at age 70 (Binomial proportion test; p-value = 3.3 × 10^−3^). The stringent FDR cutoff for discovery ensures low proportion of false positive eGenes at each age and the liberal FDR for validation ensures low proportion of false age-specific eGenes. Results remained the same for a range of discovery and replication FDR thresholds, as well as minor allele frequency thresholds (Fig. S7B).

**Figure 3:**
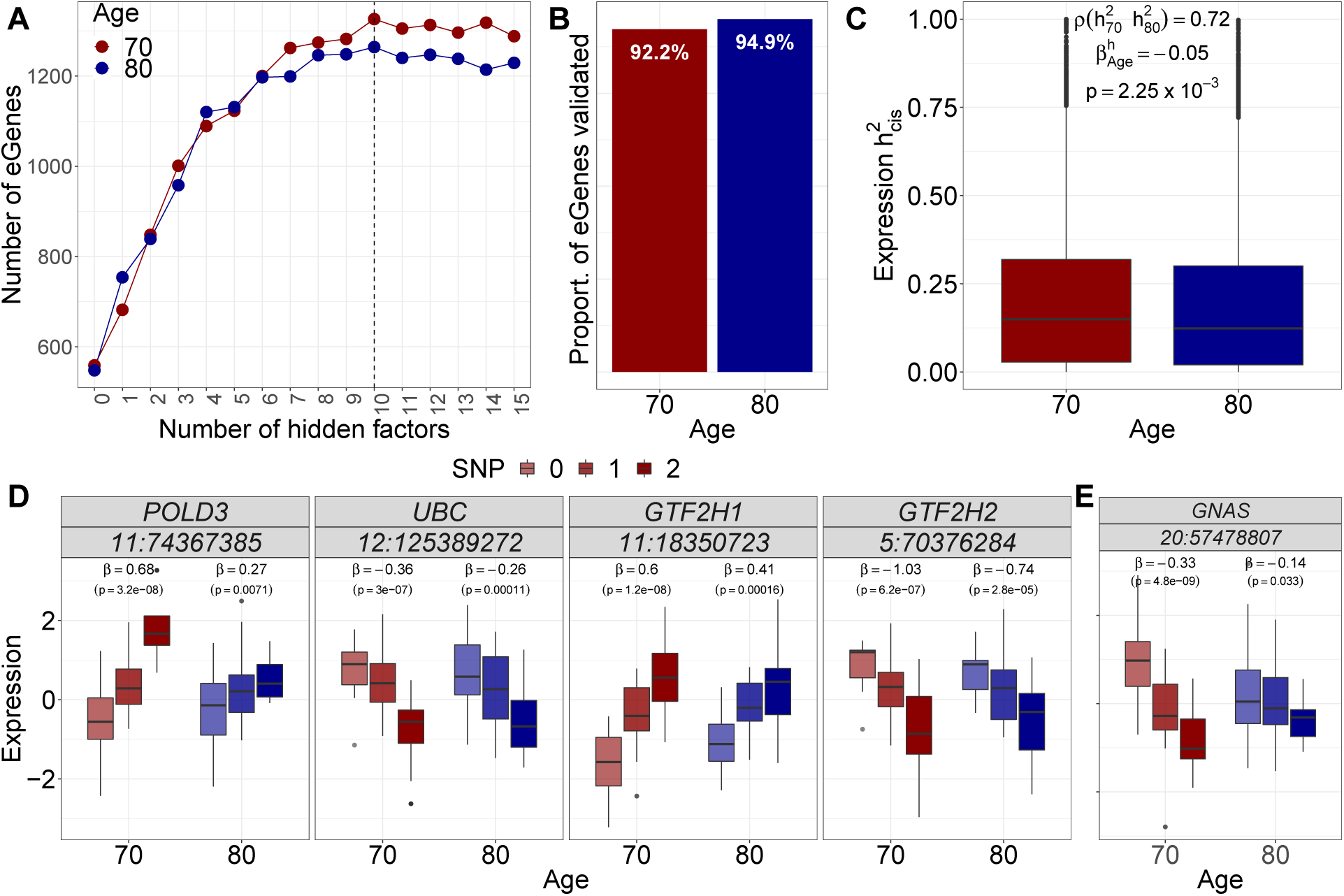
Age-specific genetic regulation across the genome. **(A)** Number of genes with at least one significant eQTL (eGenes, FDR ≤1%) for uncorrected analysis (number of hidden factors = 0) and analysis corrected for one up to 15 hidden factors. We detected 1,326 and 1,264 eGenes at age 70 and 80 (FDR ≤ 5%), respectively. The depletion of eGenes at age 80, relative to age 70, is statistically significant (Exact McNemar’s test; p-value = 5.8 × 10^−3^). Dashed line indicates the number of hidden factors that maximizes discovery at each age, i.e. 10 for both age 70 and 80. **(B)** Proportion of eGenes discovered (FDR ≤1%) at age 70 (80) that validated (FDR ≤10%) at age 80 (70). The validation proportion of eGenes discovered at age 70 is significantly smaller than the proportion at age 80 (Binomial proportion test; p-value = 3.3 × 10^−3^), indicating a decrease in genetic regulation with age. **(C)** Expression cis-heritability at each age for each gene. The decrease in average cis-heritability with age is statistically significant (average heritability at age 70 and 80 was 0.18 and 0.17, respectively; p-value = 2.25 ×10^−3^). **(D)** Genes involved in the DNA repair pathways**^?^** show loss of genetic regulation with age. For each gene, box-plot shows the expression as a function of increasing number of SNP reference alleles (color intensity) at age 70 (red) and 80 (blue) for the SNP with the largest change in effect size between the two ages; *β* and *p* -*value* indicate the eQTL effect size and p-value as estimated from MatrixEQTL. **(E)** *GNAS*, a known marker of clonal hematopoiesis**^? ?^**, is among the genes with the largest loss in genetic regulation with age. Box-plot colors are same as D.

*POLD3*, a gene with an important role in genome stability**^?^**, is among the genes with the largest loss in genetic regulation with age (Fig. 3D). Three more genes involved in the nucleotide excision DNA repair pathway, i.e. *UBC*, *GTF2H1*, and *GTF2H2*, also showed loss of genetic regulation with age; nucleotide excision DNA repair has been shown to mitigate the adverse biological effects of UV light in the exposed skin**^?^** and is associated with age-related vascular dysfunction**^?^**. In addition, *GNAS*, a key component of many signal transduction pathways and a known marker of age-related clonal hematopoiesis**^? ?^**, also showed one of the largest losses of genetic regulation with age (Fig. 3E). *GTF2H1* is the only gene mentioned above that was among the 10 genes which showed both a loss of genetic regulation and DE with age. Moreover, only 2% of the genes that showed loss in genetic regulation with age are signature genes with cell-type specific expression, indicating that our observations are very unlikely to be driven by differences in cell-type composition with age.

We estimated the *cis* heritability for each gene, i.e. the proportion of expression variance explained by *cis* SNPs, at each age using bi-variate linear mixed models (Methods; Fig. 3C). Consistent with results above, we found a small but statistically significant decrease in average *cis* heritability with age (Wald test; p-value = 2.25 × 10^−3^); the average heritability decreased from 18% at age 70 to 17% at age 80. We also estimated the genetic correlation of expression between the two ages, i.e. the proportion of expression variance shared between ages due to genetic causes. We observed a high genetic correlation of expression (*ρ*_*G*_ = 0.96; median across genes) and a strong correlation of the fixed effect sizes between the two ages (*ρ*_*β*_ = 0.70).

### 2.4. Allele-specific expression by age across the transcriptome

We investigated transcriptome-wide patterns of allele-specific expression (ASE) with age. At the global level, we found a moderate correlation of allelic ratios within an individual (Spearman’s *ρ* = 0.57; Fig. 4A). Moreover, we observed a 2.6% increase in allelic imbalance (AI) with age (median across individuals and sites; Wilcoxon signed rank test; p-value = 1.6 × 10^−2^; Fig. 4B). At the local level, as with total expression, we focused on both population-and individual-level differences in ASE with age. The former analysis requires the sites to be heterozygous across multiple individuals and captures, among others, age-interacting cis-regulatory effects while the latter captures effects of rare/personal variants or somatic mutations, e.g. as a result of clonal hematopoiesis.

**Figure 4:**
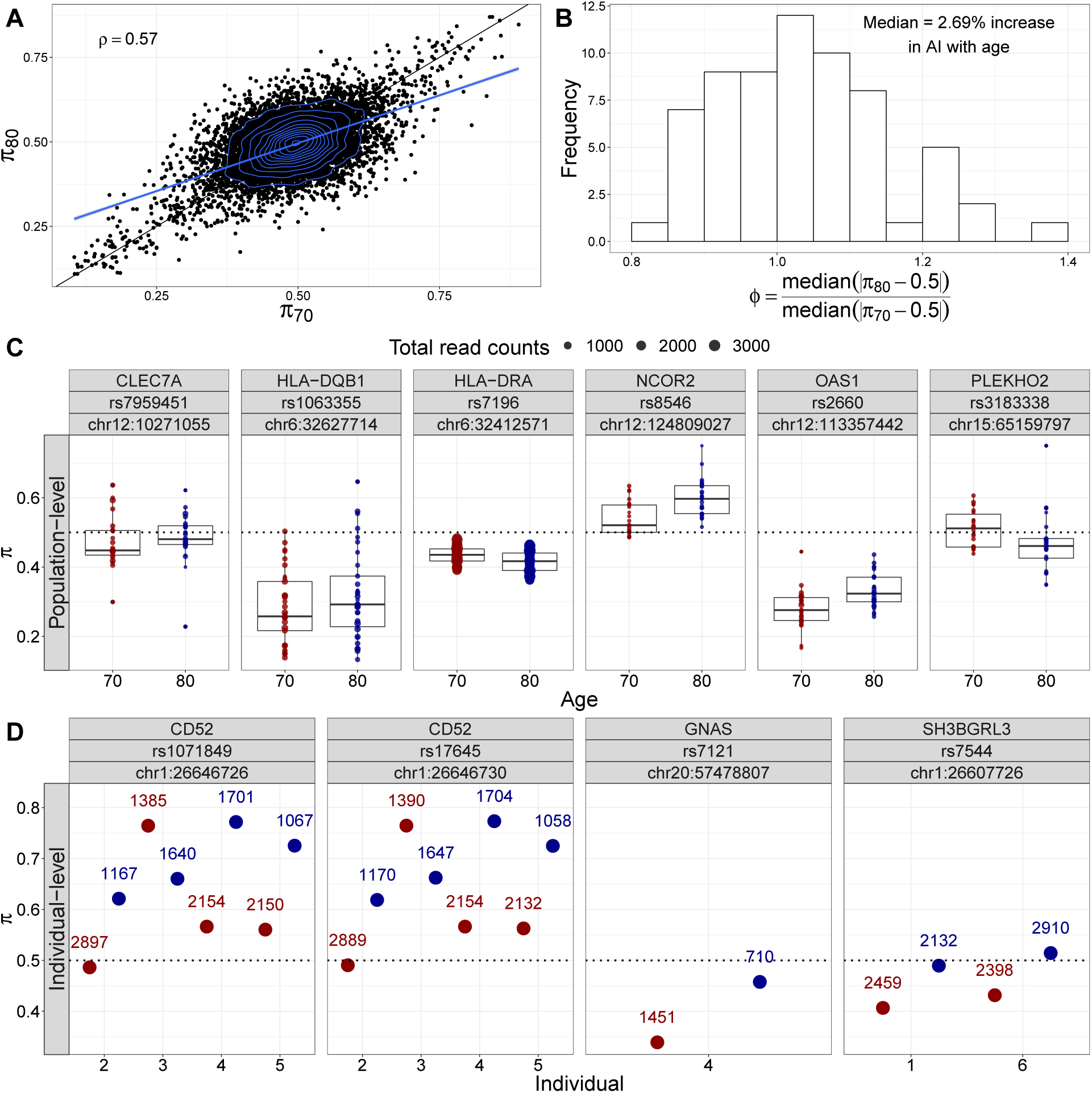
Population-and individual-level allele-specific expression across the genome. **(A)** Allelic ratios (*π*) at the two ages of each individual are moderately correlated (Spearman’s *ρ* = 0.57, across individuals and sites). Blue line shows linear regression fit while blue contour shows the 2-D density plot. **(B)** Global allelic imbalance increases with age by 2.69% (median across individuals and sites; Wilcoxon signed rank test; p-value = 1.6 × 10^−2^). Distribution of median allelic imbalance *ϕ* = | *π* 0.50|) ratio of the two samples of each individual across all heterozygous sites. **(C)** Sites with significant local population-level differential ASE with age effects. Six sites show significant differential ASE with age effects. **(D)** Six individuals on three genes, i.e. *CD52*, *GNAS*, and *SH3BGRL3*, and four sites show individual-level differential ASE with age (FDR ≤ 5%). Points indicate allelic ratio for each individual and site; color indicates age. Numbers on top of points show the total number of reads supporting the site at each age.

At the population level, sites from six genes showed significant differential ASE with age (FDR≤ 5%; Fig. 4C). *HLA-DRA*, *NCOR2*, and *PLEKHO2* show a significant gain of AI with age (Likelihood Ratio Test (LRL); p-values=9.7 × 10^−6^, 4.1 × 10^−5^, and 4.2 × 10^−5^, respectively) while *CLEC7A*, *OAS1*, and *HLA-DQB1* show a significant loss of AI with age (LRT; *p* – *values* = 7.7 × 10^−6^, 4.1 × 10^−5^, and 3.9 × 10^−6^, respectively). Most of these genes are involved in the immune system and have been previously implicated in the aging process**^? ?^**. Most notably, *NCOR2* expression and its occupancy on peroxisome proliferator-activated receptor (PPAR) target gene promoters are increased with age in major metabolic tissues. Shifting its repressive activity towards PPARs, by selectively disabling one of its two major receptor-interacting domains, resulted in premature aging in mice and related metabolic diseases accompanied by reduced mitochondrial function and antioxidant gene expression**^?^**. *CLEC7A* is the only gene with differential ASE that is also significantly down-regulated with age and a signature gene with macrophage-specific expression.

At the individual level, six individuals show differential ASE with age in four sites from three genes (FDR*<* 5%; Fig. 4D). Most notably, *GNAS*, which showed a large loss of genetic regulation with age and a nominally significant loss in population-level AI with age (LRT p-value=3.4 × 10^−3^), also showed a significant decrease in individual-level AI with age. Moreover, two individuals showed a significant loss in AI with age for *SH3BGRL3*, a gene whose expression mean has been shown to decrease with age in skin**^?^** and whose expression variance has been shown to increase with age in rat retina**^?^**. None of these genes showed significant DE with age or known to have cell-type specific expression.

### 2.5. Age-specific alternative splicing across the transcriptome

We investigated associations of transcriptome-wide changes in alternative splicing with age and discovered 503 clusters of alternatively excised introns from 294 genes (Methods) whose splicing levels were significantly associated with age (FDR ≤ 5%, Fig. 5A, Tab. S9); 11% of these genes showed also significant DE with age. GO enrichment analysis showed significant enrichment (FDR≤ 5%) of terms related to regulation of RNA splicing, apoptosis, and leukocyte differentiation (Tab. S10). The strongest associations with age were found for genes related to the circadian rhythm (Fig. 5B), disruption of which accelerates aging**^?^**, i.e. *SFPQ*, *PER1*, and *SETX*.

**Figure 5:**
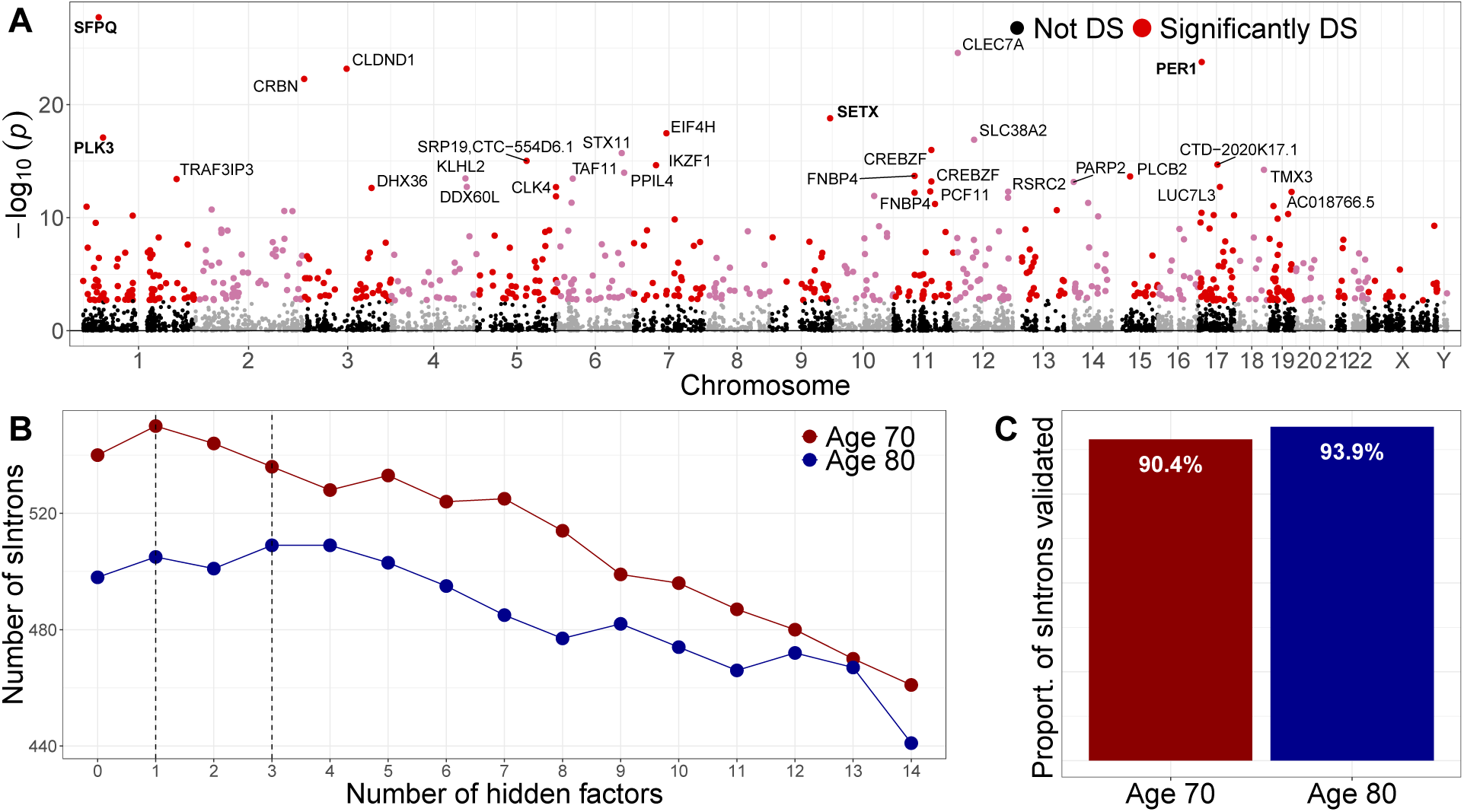
Population-level age-specific splicing across the transcriptome. **(A)** Manhattan plot of the splicing-age discoveries. Each dot represents a cluster in a gene; the x-axis gives the position of the cluster in the genome and the y-axis represents the strength of the associationwith age. We find 294 age-associated genes with 503 clusters of alternatively excised introns (3.4% of tested genes and clusters; FDR ≤ 5%). Three of the top ten genes that had the strongest association with age, i.e. *SFPQ*, *PER1*, and *SETX*, are related to the circadian rhythm, disruption of which accelerates aging**^?^**. *PLK3*, which was also in the top ten most associated genes, is implicated in stress responses and double-strand break repair. **(B)** Number of introns with at least one significant sQTL (sIntrons) at age 70 and 80 for uncorrected analysis (number of hidden factors = 0) and analysis corrected for one up to 14 hidden factors. We detected 550 and 509 sIntrons at age 70 and 80 (FDR ≤ 5%), respectively. The depletion of sIntrons at age 80, relative to age 70, is statistically significant (Exact McNemar’s test; p-value = 8.6 ×10^−3^). Dashed line indicates the number of hidden factors that maximizes discovery at each age, i.e. one and three at age 70 and age 80. **(C)** Proportion of sIntrons discovered at age 70 (80) that validated (FDR ≤ 20%) at age 80 (70). The validation proportion of sIntrons at age 70 is significantly smaller than the proportion at age 80 (Binomial proportion test; p-value = 2.2 × 10^−2^), indicating a decrease in genetic regulation with age.

*SFPQ* regulates the circadian clock by repressing the transcriptional activator activity of the *CLOCK-ARNTL* heterodimer and plays a role in the regulation of DNA virus-mediated innate immune response. Intron retention and nuclear loss of *SFPQ* are molecular hallmarks of Amyotrophic Lateral Sclerosis (ALS)**^?^**. *PER1* is a member of the Period family of genes and is expressed in a circadian pattern in the suprachiasmatic nucleus, the primary circadian pacemaker in the mammalian brain. Genes in this family encode components of the circadian rhythms of locomotor activity, metabolism, and behavior. This gene is upregulated by *CLOCK-ARNTL* heterodimers, but then represses this upregulation in a feedback loop using PER/CRY heterodimers to interact with *CLOCK-ARNTL*. *SETX* is implicated in transcription termination and DNA double-strand breaks damage response generated by oxidative stress**^?^**. Mutations in this gene have been associated with juvenile ALS**^?^**. *SETX* is also required for the transcriptional termination of PER1 and CRY2, thus playing an important role in the circadian rhythm regulation.

### 2.6. Age-specific genetic regulation of alternative splicing across the genome

We tested for genetic variants that affect alternative gene splicing on the autosomes. For each gene, we quantified intron usages with LeafCutter**^?^** and evaluated the association between intron usage ratios at each age and genetic variants within 100kb of the intron (Methods). After correction for background noise (Fig. 5C), we detected significant sQTLs for 550 introns at age 70 (1.4% of tested introns, FDR ≤ 5%, Tab. S11), 90.4% of which replicated at age 80 (FDR ≤ 20%, Fig. 5D). On the other hand, we detected sQTLs for 509 introns at age 80, 93.9% of which replicated at age 70. The depletion of introns with at least one significant sQTL (sIntrons) at age 80, relative to age 70, was statistically significant (Exact McNemar’s test; p-value = 8.6 × 10^−3^). Moreover, the proportion of sIntrons discovered at age 70 that replicated at age 80 was significantly smaller than the proportion of sIntrons discovered at age 80 and replicated at age 70 (Binomial proportion test; p-value = 2.2 × 10^−2^), indicating a small but significant loss of sQTLs as individuals age.

## 3. Discussion

We have studied the combined effects of age and genetics on gene expression and alternative splicing in 65 humans from the general population sampled twice ten years apart. Our focus on 70-and 80-year old elderly individuals was designed to capture transcriptome changes during a period of high morbidity and mortality; the average life expectancy in Sweden is 80 and 83 years for men and women, respectively. We observed that individuals were more similar to their own gene expression levels between the two ages than to other individuals of the same age. This indicates that a larger proportion of gene expression variance is explained by shared genetics and environment than by the advanced aging process.

Despite the relative stability of expression profiles within individuals over time, we were able to identify 1,291 genes with significant changes in expression. Pathways related to the adaptive immune system, cell signaling, and inflammatory response were among those enriched for down-regulated genes while up-regulated genes were enriched for pathways related to oxidative phosphorylation, adaptive immune system, and metabolism of proteins. Many of these functions have been previously described as hallmarks that represent common denominators of aging**^?^**. Moreover, 18 of the differentially expressed genes are previously known to be complex-trait associated genes where gene expression levels modulate risk.

Because the rate of aging varies among individuals, humans become increasingly different from each other with age**^?^**. Thus, chronological age fails to provide an accurate indicator of the aging process. Longitudinal studies offer a better understanding of the aging process by studying the same individuals throughout their lifespan, collecting serial assessments rather than by comparing individuals of different ages from different environments. Recent longitudinal studies have begun to highlight the importance of individual-level molecular profiling to identify important health factors**^? ?^**. Using our longitudinal design, we were able to analyze individual changes in gene expression and identified 529 individually-dynamic genes with functions related to regulation of proteolysis and immune response. The sharing of immune-related function for both population and individual differentially-expressed genes may, in part, be explained by observations of increased immune dysregulation and transcriptional variability with age**^? ?^**.

Genetic regulation of gene expression is involved in the etiology of many complex human traits**^? ?^**. Previous studies in model organisms have reported a reduction in these associations with age**^?^**; however, less is known about the extent of genetic dysregulation with age in humans. Cross-sectional studies in humans have identified age-specific eQTLs for three**^?^** and ten**^?^** genes, respectively. A smaller two-year longitudinal study of middle-aged females found two genes with time-dependent associations**^?^**. Here, we report a global reduction in genetic control and a reduction in gene expression heritability with age. This reduction could be due to several factors, such as the diminution of the level of expression of transcription factors, epigenetic modification, or genomic instability. Notably, while aging led to a reduction in genetic control, we observed an increase in the levels of allele-specific expression with age. The high correlation of genetic effects and the increase in allelic imbalance with age suggests that increasing environmental variance as opposed to decreased genetic variance underlies a component of the reduction in heritability and loss of genetic effects.

Deregulation of precursor mRNA splicing is associated with many illnesses and has been linked to age-related chronic diseases. There are no prior longitudinal studies of the human transcriptome**^? ?^** assessing the dynamics of alternative splicing and its genetics. We found 294 genes, related, among others, to regulation of RNA splicing and apoptosis, with age-specific alternative splicing. Three of the top ten genes with the strongest association of alternative splicing with age are related to the circadian rhythm, disruption of which is known to accelerate aging**^?^**. In addition to changes in alternative splicing with age, we also observed a reduction in the number of genetic associations with splicing between the two ages highlighting similar patterns of dysregulation for both expression and splicing.

In addition to immune response and circadian rhythm, we observed that genes that lose regulation with age are involved in DNA repair pathways. For example, *POLD3*, a gene with an important role in genome stability, was among the genes with largest loss of genetic control with age. In addition, *GNAS*, a known marker of clonal expansion, was also among the genes with the largest loss of genetic control with age and showed a significant decrease in allele specific expression with age. Moreover, *PLK3*, a gene implicated in stress responses and double-strand break repair, was among the top ten genes with the strongest age-alternative splicing association.

In summary, we present the first long-term, longitudinal characterization of expression and splicing changes as a function of age and genetics. Our findings indicate that, although gene expression and alternative splicing and their genetic regulation are mostly stable late in life, a small subset of genes is dynamic and is characterized by changes in expression and splicing and a reduction in genetic regulation, most likely due to an increase of environmental variance and de-regulation of DNA repair pathways.

## 4 Materials and Methods

### 4.1. Study cohort

The Prospective Investigation of Uppsala Seniors (PIVUS) study is a population-based study of the cardiovascular health in the elderly**^?^**. The PIVUS cohort is comprised of 1,016 individuals (509 females and 507 males) of Swedish ancestry living in Uppsala, Sweden from 2001 to 2005. Participants were examined at age 70 and 80 with deep phenotyping and blood was frozen upon collection. Our focus on 70-and 80-years old elderly individuals was designed to capture changes during a period of high morbidity and mortality; the average life expectancy in Sweden is 80 and 83 years for men and women, respectively. A detailed description of the recruitment and phenotype data for this cohort is provided elsewhere**^?^**. The Ethics Committee of the University of Uppsala approved the study, and the participants gave their informed consent.

### 4.2. RNA isolation and sequencing

Gene expression was quantified for 65 individuals at both ages (130 samples). Total RNA was extracted from 400 *µ*L whole blood using the NucleoSpin RNA Blood Kit (Macherey-Nagel, Düren, Germany) according to the manufacturer’s directions. Samples were eluted in 60 *µ*L RNase-free H_2_O. A small aliquot of each sample was set aside for quality assessment and the remainder was immediately stored at −80C. The RNA yield was estimated by measuring absorbance at 260 nm on the Nanodrop 2000 (Thermo Fisher), and RNA purity was determined by calculating 260/280 nm and 260/230 nm absorbance ratios. RNA integrity was assessed on the Agilent Bioanalyzer using the RNA 6000 Nano Chip kit (Agilent Technologies). An RNA integrity number (RIN) was assigned to each sample by the accompanying Bioanalyzer Expert 2100 software.

cDNA libraries were constructed following the Illumina TrueSeq Stranded mRNA Sample Prep Kit protocol and dual indexed. The average size and quality of each cDNA library was determined by the Agilent Bioanalyzer 2100 and concentrations were determined by Qubit for proper dilutions and balancing across samples. Twenty pooled samples with individual indices were run on an Illumina NextSeq 500 (high output cartridge) as 2×75 paired end sequencing. Output BCL files were FASTQ-converted and demultiplexed.

### 4.3. Genotyping and imputation

DNA was extracted and genotyped on the Illumina OmniExpress and Cardio-Metabochip arrays for more than 700K SNPs. Genotype data quality control was described elsewhere**^?^**. Only 63 of the 65 RNA-Sequenced individuals passed genotype quality control thus all analyses involving genotype data, e.g. eQTL analysis, were performed on these 63 individuals with complete data. Genotype data was phased using SHAPEIT**^?^** and imputed with Impute2 (v2.3.2)**^?^** using the CEU haplotypes from the 1000 Genomes Project Phase-3 reference panel**^?^**. Post imputation quality control is described in the Supplemental Material.

### 4.4. RNA-Seq quality control

Picard, Samtools**^?^**, and other metrics were used to evaluate data quality (Fig. S1). Only genes that passed expression threshold were used; genes were considered expressed if, at both ages, they had, on average, at least 5 counts and zero counts in no more than 20% of individuals (to minimize tails). A total of 16,086 genes were considered expressed. Gene expression data was library-size-corrected, variance-stabilized, and log2-transformed using the R package DESeq2**^?^**. We refer to this version of the data as ‘raw data’ as it is not corrected for global determinants of gene expression variability (see below).

### 4.5. Background noise correction of gene expression data

In order to identify and correct for major components of gene expression variability, unrelated to age, we used surrogate variable analysis (SVA) as implemented in the R package `smartSVA`**^?^**, setting age as variable of interest (Fig. S3). More details about the implementation of SVA can be found in the Supplemental Methods. We selected SVA, as opposed to other background noise correction methods**^? ?^**, on the basis of recent work that shows it is robust to spurious associations when setting a variable of interest**^?^**. Using the **?** method, we estimated the number of hidden factors that explains a significant amount of the expression variability to be 15.

All results in the main paper are corrected for hidden factors extracted by SVA as well as RIN and RNA concentration, two technical covariates moderately correlated with age (Fig. S3) that could act as potential confounders, to avoid false positives. In the Supplement, we show results from uncorrected analyses and analyses corrected for measured factors (Table S1), or hidden factors extracted by SVA without setting age as a variable of interest.

### 4.6. Hierarchical clustering of gene expression

We performed hierarchical cluster analysis on the sample-to-sample distance matrix of the expression data. To compute the sample-to-sample distance matrix, we used the R function hclust from the stats package. We used the Euclidean distance measure to determine the distance between sets of observations. We used the *complete linkage* clustering strategy, a method that aims to find similar clusters. Samples were classified as ‘clustered with individual ID’ if their nearest neighbor based on the dendrogram was their own sample from another age.

### 4.7. Differential expression analysis

For each gene, we fit the following linear mixed model: expression ∼ individual (random) + age (fixed) + 15 hidden factors (fixed) + RIN (fixed) + RNA concentration (fixed), using the lme4 R package**^?^**. Age was coded as 0 and 1 for individuals at age 70 and 80, respectively. P-values were calculated based on Satterthwaite’s approximations implemented in the lmerTest R package**^?^**. Significance of the results was assessed using the **?** q-value method implemented in the qvalue R package to control the FDR at 5%.

### 4.8. Enrichment analysis for differentially expressed genes

Pathway enrichment analyses for DE genes were performed using GSEA**^?^**, a computational method that determines whether an a priori defined set of genes shows statistically significant, concordant differences between two biological states (here age 70 vs 80). Resulting p-values are adjusted for multiple testing using the `q-value` method**^?^** controlling FDR at 5%.

### 4.9. Replication of differential expression results in other blood studies

We assess the significance of the overlap between our top 1,000 DE genes from PIVUS and the top 1,000 DE gene in CHARGE**^?^** and SardiNIA**^?^** using the exact test of multiset intersection implemented in the R package `SuperExactTest`**^?^**. We perform GO enrichment analysis for intersect genes that are shared between the three studies using WebGestaltR**^?^**.

### 4.10. Identification of individuals with outlying age trajectories

We only looked for outliers on autosomal genes and among the 61 individuals that cluster with their own sample at another age (Fig. 1A). Individuals are outliers for a gene, if their change in the expression of the gene between the two ages (*E*_80_ − *E*_70_) falls outside the (*Q*_1_ − 3 × *IQR, Q*_3_ +3 × *IQR*) range, where *Q*_1_ and *Q*_3_ are the 25th and 75th percentiles and IQR is the interquartile range of the distribution.

### 4.11. eQTL mapping

We mapped eQTLs at each age using the linear regression models implemented in the `MatrixEQTL` R package**^?^**. For the analysis correcting for measured/hidden factors, we include the measured/hidden factors as covariates. To call eGenes at each age, i.e. genes with at least one significant eQTL, as well as eQTLs for each eGene, we use the R package `TreeQTL`**^?^** controlling the FDR at 1%, both at the gene and gene-SNP level. The final number and list of eGenes at each age was obtained using the number of expression hidden factors that maximized discovery at each age, i.e. 10 hidden factors (Fig. 3A).

### 4.12. Age-specific eQTL mapping

In order to identify eGenes that are specific to age 70, i.e. show loss of genetic regulation with age, we use a two-step FDR approach from validation theory**^?^**. Specifically, we first discover eGenes at age 70 at 1% FDR, as described above, and then we validate them at age 80 at 10% and 20% FDR, by performing eQTL mapping only for these genes. In order to identify eGenes that are specific to age 80, we reverse the process.

### 4.13. Heritability of gene expression at each age

For each gene, we used the bivariate GREML method, implemented in the GCTA software**^?^**, to estimate the cis-heritability of expression at each age as well as the genetic correlation of expression between the two ages. To estimate the average cis-heritability with age, we use the Beta regression models implemented in the R package `betareg`**^?^**, modelling the logit of the cis-heritability of each gene as a function of age, i.e. *logit*(*h*^2^) = *α* +*β* × *age*, where *h*^2^ is the estimated cis-heritability of a gene, *β* is the effect of age on heritability, and *age* is coded as above. Then, the estimated average cis-heritability at age 70 and 80 is given by 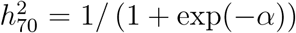 and 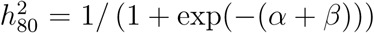 respectively. When testing for the significance of the difference in cis-heritability with age, we also adjust for the standard error of the heritability, in order to take into account cases where the estimate of the heritability at one of the two ages is noisier.

### 4.14. Quantifying allele-specific count data from RNA-seq data

Read mapping bias was removed by following the WASP pipeline**^?^**. The GATK tool `ASEReadCounter` was used to count reads at exonic heterozygous sites. Only bi-allelic SNPs with allelic ratio ≥.1 or ≤.9 were considered. Correlation of allelic ratios between the two ages and testing for changes in global AI and local individual-level differential ASE with age was performed using only sites supported by at least 50 reads at each age of an individual. The local population-level differential ASE with age analysis was performed on sites supported by at least 20 reads at each age of an individual in at least 20 individuals.

### 4.15. Age-specific ASE mapping

Let *π*_*ijk*_ and *ϕ*_*ijk*_ = |*π*_*ijk*_ −0.5 | denote the proportion of reads supporting the reference allele and the the allelic imbalance, i.e. absolute deviation from allelic balance, for individual *i* (*i* = 1, …, 65), at age *j* (*j* = 70, 80), and heterozygous site *k* (*k* = 1*, …, K*_*ij*_). Moreover, let *ϕ*_*ij*\*_ the median allelic imbalance for *i* at age *j* and 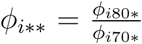 the ratio of median AI between the two ages of individual *i* across all sites. We test for a statistically significant increase in global AI with age (*H*_0_ : *µ* ≤ 1 vs *H*_1_ : *µ >* 1, where *µ* median of the *ϕ*_*i*\*\*_’s) using the one-sample Wilcoxon signed rank test.

At the local level, we test for population-and individual-level differences in ASE with age using the beta-binomial generalized linear mixed models implemented in `EAGLE` (v2.0)**^? ?^**.

### 4.16. Quantification of alternative splicing

To quantify alternative splicing events we followed the `LeafCutter`**^?^** pipeline. In short, we first mapped the 130 RNA-Seq samples from PIVUS to the human genome (hg19) using STAR, allowing de-novo splice junction predictions. We then used `LeafCutter`**^?^** to identify alternatively excised introns by pooling all junction reads. LeafCutter then defines ‘clusters’ of alternatively excised introns that represent alternative splicing choices. This resulted in 78,373 alternatively excised introns from 24,126 clusters. Each cluster comprises of, on average, 3.2 introns (median = 3, min=2, max=51).

To identify alternative splicing events that are suitable for differential splicing and sQTL by age analysis, we first exclude clusters with more than 10 introns. Then, we restrict our analyses to active introns, i.e. introns that are supported by at least 10% of the total number of reads supporting the clusters they belong to in at least 25% of samples, considering each age separately. Clusters with less than two active introns after this step were filtered out. Last, we only consider clusters that exhibit some minimum splicing variability, i.e. clusters with Hellinger’s distance ≥ 1%**^?^**. In the end, we have 14,917 clusters and 36,713 introns.

### 4.17. Background noise correction of alternative splicing data

As with the gene expression data, we used SVA**^?^**, setting age as variable of interest, to identify major components of alternative splicing variability (Fig. S8). More details about the implementation of SVA can be found in the Supplemental Methods. Using the **?** method, we estimated the number of hidden factors that explains a significant amount of the alternative splicing variability to be 14.

### 4.18. Differential splicing by age analysis

To identify alternative splicing events with age, we used the Dirichlet-multinomial generalized linear model implemented in `LeafCutter`**^?^**. We adjust our analysis for measured factors that explain more than 0.5% splicing variability, i.e. RIN, proportion of intronic bases, median insert size, and extraction year (Fig. S8B). P-values of association with age were calculated based on the likelihood ratio test. To call dsGenes, i.e. genes with at least one significantly DS cluster, as well as DS clusters for each dsGene, we use the R package `TreeQTL`**^?^** controlling the FDR at 5% both at the gene and gene-cluster level.

### 4.19. sQTL mapping

We used linear regression, as implemented in Matrix eQTL**^?^**, to test for associations between ratios of alternatively excised introns at each age group and variants within 100kb of the intron clusters, adjusting for splicing hidden factors. We control the FDR at 5% both at the intron and intron-SNP level using `TreeQTL`**^?^**.

## Data Access

fastq files containing sequence data have been deposited in the European Genome-phenome Archive (EGA-box-1167).

## Acknowledgments

Research leading to this manuscript was supported by the Stanford Genome Training Program, the Stanford University School of Medicine Dean’s Postdoctoral Fellowship, the Glenn Institute for Aging at Stanford, the Stanford Graduate Fellowship in Science and Engineering, and NIH R01 grant DK106236.

## Author Contributions

Conceptualization, E.I. and S.B.M.; Methodology, B.B., S.J., C.S., E.I., and S.B.M; Formal Analysis, B.B., M.D., O.d.G, N.S.A., X.L, B.L, and M.J.G.; Investigation, N.L.C. and K.S.S.; Resources, M.P., F.C., D.S., L.L, N.L.C., and K.S.S.; Writing-Original Draft, B.B., E.I., and S.B.M; Writing-Review & Editing, B.B., M.D., O.d.G, N.S.A., X.L, B.L, M.J.G., N.L.C., D.S., C.S., L.L., E.I., and S.B.M;Funding Acquisition, E.I. and S.B.M; Supervision, E.I. and S.B.M.

## Supplemental Material

### S1. SNP array data and imputation

Genotype data quality control was described elsewhere**^?^**. In summary, 949 individuals passed genotype quality control. Genotype phasing and imputation was performed in all 949 individuals that passed quality control. Post-imputation quality control was performed as follows. SNPs with an imputation info-score below 0.4, a HWE P-value ≤ 10^−6^, or a MAF ≤ 5% in the 63 individuals with measured RNA-Seq and methylation were excluded. In total, 7,037,776 SNPs passed post-imputation quality control.

### S2. Expression quantification and quality control

The quality of the raw reads was assessed using FastQC (v0.11.5). The adaptors were clipped using cutadapt (v1.8.1)**^?^** requiring at least three bases to match (–min overlap 3) and removing processed reads shorter than 20 bases (–min length 20). RNA-Seq reads were mapped to the NCBI v37 H. sapiens reference genome using STAR (v2.4.2a)**^?^**. Only uniquely aligned reads were used for downstream quantification and analysis. The percentage of reads marked as PCR duplicates was computed using Picard. For the differential expression and eQTL analysis, PCR duplicates were not filtered out, since it has been shown that computational removal of duplicates does not improve power or FDR in differential expression analyses**^?^**, but the proportion of PCR duplicates used as a technical covariate in downstream analysis. For the differential ASE PCR duplicates were removed. Mapping statistics from the BAM files were acquired through Samtools flagstat (v1.2)**^?^**. The 5’ and 3’ coverage bias, duplication rate and insert sizes were assessed using Picard tools (v2.0.1). HTSeq was used to quantify gene expression**^?^**.

Expression data on the sample level were first corrected for library size using the `DESeq2` R package**^?^**. Genes were excluded if they had less than 5 counts on average for either age groups, and zero counts in more than 20% of individuals (to minimize tails). A total of 16,087 genes were expressed. For the eQTL analysis, genes from the sex chromosomes as well as mitochondrial genes were also excluded, leaving 15,729 genes in the analysis.

To identify potential outlier individuals, we performed PCA based on total and decomposed gene expression measurements (see Methods below; Figure S1). Samples that demonstrated extreme values in the first two principal components on the expression levels were removed (more than 3 standard deviations). No samples were excluded based on this measure. To identify low quality samples, we applied several quality metrics (Figure S1). Samples were removed if they had insufficient reads (≤20M), poor mappability (≤60%), and low correlation with other samples (D-statistic ≤ 0.85). We also checked for sample mix-ups by comparing agreement between true and RNA-Seq-inferred heterozygous SNPs for all possible pairs of RNA-Seq and genotype data. No samples were excluded based on these metrics. Three individuals with low RNA integrity number (RIN ≤ 5) were marked. These individuals were not excluded since they do not seem to appear as outliers in the PCA plots (Figure S1C).

### S3. Measuring DNA methylation and quality control

Extracted DNA was bisulphite-converted using the Zymo bisulphite conversion kit and hybridized to the Infinium HumanMethylation450 BeadChip (450k array). Signal intensities were measured with the BeadChip scanner. The 130 samples (with available RNA-Seq data) were distributed across two periods of methylation data collection, eight 96-well plates, and forty-one 12-sample chips.

Quality control was performed using the R package `minfi` (version 1.20.0)**^?^**. Missing values were imputed by the k-nearest neighbor approach using the impute R package (ver-sion 1.48.0)**^?^**.Background correction and dye-bias normalization were done using noob**^?^** through minfi. Signal intensities were converted to DNA methylation *β*-values, i.e. the ratio of methylated probe intensity to total (methylated + un-methylated) probe intensity.

### S4. Inferring cell-type frequencies from whole blood

Whole blood is a heterogeneous mixture of cell types. Since gene expression and DNA methylation vary across different cell types, correlations between the phenotype of interest (e.g. age) and the cell type composition may lead to a large number of false discoveries. False discoveries due to cell type heterogeneity can be addressed by adding the cell proportions as covariates. Since cell counts were not available for our samples, we used computational methods to estimate their composition.

To estimate blood cell composition from gene expression data, we used CIBERSORT**^?^**. While this tool was designed from micro-array data, it has been shown to have reasonably robust cross-platform performance. To estimate blood cell composition from DNA methylation data, we used Houseman’s reference-based method**^?^**, including their provided reference data and signature CpG sites. This approach was used on our methylation data after adjustment with `noob`**^?^**. Results are shown in Figure S2A.

Methylation-based cell-type frequency estimates are closer to expected values for adults of similar age, compared to expression-based estimates. Moreover, while methylation-based estimates are mainly correlated with biological covariates, expression-based estimates are highly correlated with technical factors Figure S2B. For these reason, we use methylation-based estimates in downstream analyses. Methylation-based estimates of B and CD8 T cells, granulocyte, and monocytes showed a significant difference between the two ages (2-sample Wilcoxon test; p-values = 0.018, 0.019, 1.21 × 10^−4^, and 9.29 × 10^−3^, Figure S2C).

### S5. Background noise correction in RNA-Seq experiments

We consider analyses corrected for either measured and/or inferred determinants of gene expression variability. Below we describe the selection of the known factors and the inference of the inferred factors.

#### S5.1. Measured determinants of gene expression variability in RNA-Seq experiments

We considered 24 measured variables as candidate components of RNA-Seq variability, listed in Table S1. In order to decide which of the variables affect gene expression, we performed a multiple linear mixed model regression on the expression of each gene using the lme4 R package**^?^**. We used the *π*_1_ statistic**^?^** to detect technical covariates affecting a large number of genes, i.e. *π*_1_ ≤ 5%, and only consider those covariates in subsequent analyses. Table S1 also lists the median % of gene expression variance accounted for (VAF) by each measured variable, estimated using the R package `variancePartition`**^?^**, as well as the proportion of genes each variable was associated with at 5% FDR.

Figure S3a and c show the correlation of the variables (that have *π*_1_ ≤ 5%) with age and the proportion of gene expression variance they explain. Since age is moderately correlated with RIN (Spearman’s *ρ* = −0.46*, p* -*value* = 5.16 × 10^−8^) and RNA concentration (Spearman’s *ρ* = −0.30*, p* -*value* = 4.21 × 10^−4^) and RIN and RNA concentration are associated with gene expression, these variables could act as potential confounders and we thus include them in the model for differential expression analysis.

#### S5.2. Inferred determinants of gene expression variability in RNA-Seq experiments

We used surrogate variable analysis (SVA) to infer hidden factors from the RNA-Seq data. Two different algorithms for extracting hidden factors were considered: the two-step SVA procedure**^?^** without setting any covariate of interest, implemented in the sva R package**^?^**, and the IRW-SVA algorithm, setting age as the covariate of interest**^?^**, implemented in the `SmartSVA` R package**^?^**. The later algorithm, to which we hereafter refer as supervised SVA, attempts to protect the effect of age by identifying a subset of genes that show strong association with the underlying sources of gene expression heterogeneity but no association with age, also referred to as *negative* or *empirical* control genes.

We use the Buja and Eyuboglu method**^?^**, a permutation-based selection rule for the number-of-factors problem, to estimate the number of hidden factors that explain a significant proportion of gene expression variability, larger than what would be expected by chance. Using the SVA method, we find 12 and 15 factors when performing the unsupervised and supervised algorithms, respectively. On the other hand, for the MSVA, we find 29 (14 within and 15 between) and 42 (22 within and 20 between) factors via the unsupervised and supervised algorithms.

We found that the inferred factors summarize multiple correlated measured factors (Figure S3b) with significant contribution to variability in the RNA-sequencing data (Figure S3d). Generally, we observed that the top factors largely correspond to technical factors such as RNA extraction date, RIN scores and factors specific to RNA-seq such as percent duplicated reads and others obtained from the Picard metrics and to a much lesser extent to biological factors such as estimates of cell type frequencies.

### S6. Co-localization analysis for DE genes

For each DE gene, we obtained colocalization posterior probabilities (CLPP) between GWAS summary statistics of several complex traits and GTEx**^?^** whole blood from the *LocusCompare* database (http://locuscompare.ml:3838/). We defined any locus with CLPP ≥ 0.05 to have sufficient evidence for colocalization.

### S7. Differential expression analysis in the SardiNIA study

The SardiNIA study consists of 605 individuals (56% females, average age 57) from 195 families with measured RNA-Seq**^?^**. From a total of 19,646 genes expressed in SardiNIA, 14,847 of them are also expressed in PIVUS, using the same threshold for calling a gene expressed (see above).

To account for the family structure, we perform the differential expression analysis using the pedigree-based linear mixed-models implemented in the `coxme` R package**^?^**. Specifically, let *E*_*ij*_ and *Age*_*ij*_ denote the gene expression and age for the *j*^*th*^ member of family *i*, and **x_ij_** a *p*-dimensional vector with known / inferred determinants of gene expression for *i* = 1, …, *n* and *j* = 1, …, *n*_*i*_. Here we correct the analysis for sex. In the mixed models framework, a set of family-specific random effects **b_i_** = (**b_i1_**, **b_i2_**, …, **b_in_**)**^T^**is introduced to model the within family dependencies and *E*_*ij*_ is modeled conditional on the *n*_*i*_ × 1 random-effects vector **b_i_**, and covariate information *Age*_*ij*_ and **x_ij_** as

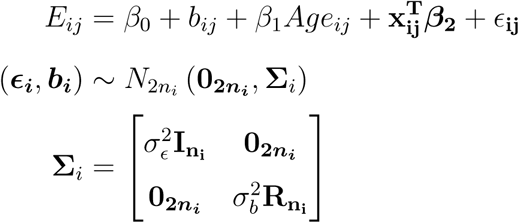

Where *β*_0_ is the intercept term, *β*_1_ is the fixed effect of age, and ***β*_2_** is the *p*-dimensional regression coefficients vector for the additional covariates. Moreover, **R_n_i** is the coefficient of relationships matrix with elements 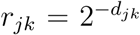 with *d*_*jk*_ denoting the distance between subjects *j* and *k* in the pedigree and 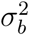 the genetic variance parameter. 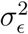 is the residual variance and **I_ni_** is an *n*_*i*_ × *n*_*i*_ identity matrix.

## Supplemental Tables

**Table S1:**
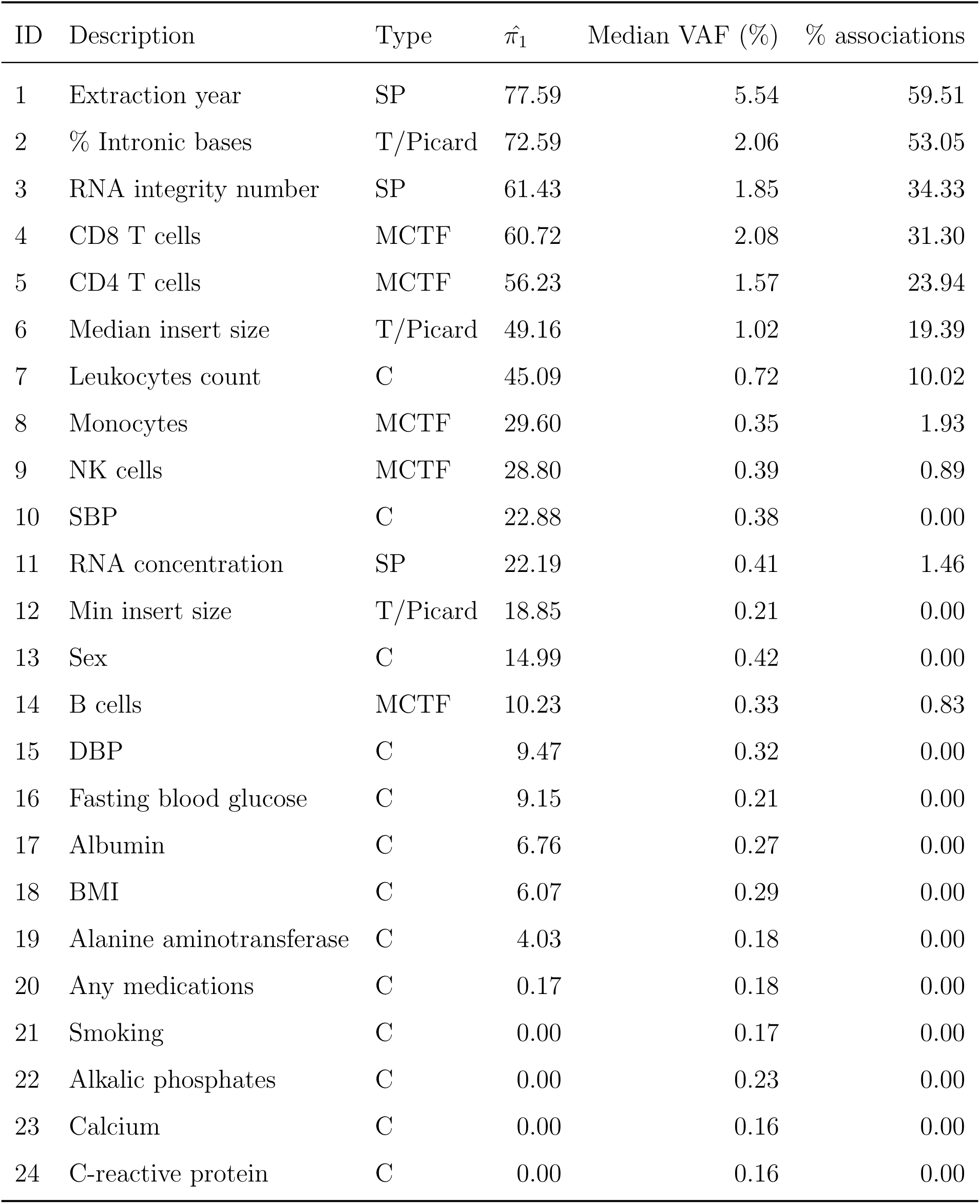
Measured covariates that can introduce variability in RNA-sequencing experiment. Table lists technical factors directly obtained from Picard QC metrics (Type “T/Picard”), factors relating to sample preparation and storage (Type “SP”), methylation-based cell type frequencies (Type “MCTF”), and factors measured in the clinic (Type “C”). For subsequent analyses, we only consider factors that affect a large proportion of genes 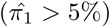. Table also shows median % of expression variance accounted for (VAF) by each factor and genes each factor was associated with at 5% FDR. 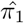, VAF, and the number of associations for each covariate were estimated via a multiple linear mixed model per gene correcting for all uncorrelated covariates. Related to Figure 1.

| ID | Description | Type | $\hat{\pi}_1$ | Median VAF (%) | % associations |
| --- | --- | --- | --- | --- | --- |
| 1 | Extraction year | SP | 77.59 | 5.54 | 59.51 |
| 2 | % Intronic bases | T/Picard | 72.59 | 2.06 | 53.05 |
| 3 | RNA integrity number | SP | 61.43 | 1.85 | 34.33 |
| 4 | CD8 T cells | MCTF | 60.72 | 2.08 | 31.30 |
| 5 | CD4 T cells | MCTF | 56.23 | 1.57 | 23.94 |
| 6 | Median insert size | T/Picard | 49.16 | 1.02 | 19.39 |
| 7 | Leukocytes count | C | 45.09 | 0.72 | 10.02 |
| 8 | Monocytes | MCTF | 29.60 | 0.35 | 1.93 |
| 9 | NK cells | MCTF | 28.80 | 0.39 | 0.89 |
| 10 | SBP | C | 22.88 | 0.38 | 0.00 |
| 11 | RNA concentration | SP | 22.19 | 0.41 | 1.46 |
| 12 | Min insert size | T/Picard | 18.85 | 0.21 | 0.00 |
| 13 | Sex | C | 14.99 | 0.42 | 0.00 |
| 14 | B cells | MCTF | 10.23 | 0.33 | 0.83 |
| 15 | DBP | C | 9.47 | 0.32 | 0.00 |
| 16 | Fasting blood glucose | C | 9.15 | 0.21 | 0.00 |
| 17 | Albumin | C | 6.76 | 0.27 | 0.00 |
| 18 | BMI | C | 6.07 | 0.29 | 0.00 |
| 19 | Alanine aminotransferase | C | 4.03 | 0.18 | 0.00 |
| 20 | Any medications | C | 0.17 | 0.18 | 0.00 |
| 21 | Smoking | C | 0.00 | 0.17 | 0.00 |
| 22 | Alkalic phosphates | C | 0.00 | 0.23 | 0.00 |
| 23 | Calcium | C | 0.00 | 0.16 | 0.00 |
| 24 | C-reactive protein | C | 0.00 | 0.16 | 0.00 |

Table S2: **Population-level age-specific expression across the transcriptome.** Summary statistics for differential expression by age analysis in the PIVUS study, provided as a separate excel file. Table lists the age effect size, s.e., p-value, BH-adjusted p-value, q-value, and proportion of variance explained for each each gene. Related to Figure 1.

Table S3: **Gene set enrichment analysis for genes differentially expressed with age.** Provided as a separate excel file. Table lists the pathway name, top level categorization (used for Figure 1), size, enrichment score (ES) as computed by GSEA, normalized ES, enrichment p-value and q-value, and number and proportion of genes in the pathway that are DE by age in PIVUS. Related to Figure 1.

Table S4: **Co-localization of GWAS traits and GTEx whole-blood eQTL summary statistics for DE genes.** The table, provided as a separate excel file, lists the DE gene names and symbols with colocalization posterior probability (CLPP) above 5% between GWAS hits and GTEx whole-blood eQTLs. Related to Figure 1.

Table S5: **Genes differential expressed by age in PIVUS, CHARGE, and SardiNIA.** Provided as a separate excel file. Table lists the gene names and their effect size sign and p-value in each of the three studies. Related to Figure 1.

Table S6: **Genes differential expressed by age in PIVUS that are known aging-and longevity-related genes from The Human Ageing Genomic Resources (HAGR) GenAge and LongevityMap databases.** Provided as a separate excel file. Table lists gene symbol and description and the HAGR database name. Related to Figure 1.

Table S7: **Individual-level age-specific expression across the transcriptome.** Summary statistics for age-trajectory outliers provided as a separate excel file. The table lists the outlier gene symbol, description, and chromosome, the outlier sample ID and Z-score, and the outlier extremeness. Related to Figure 2.

Table S8: **Age-specific genetic regulation of gene expression across the transcriptome.** Summary statistics for the eQTL analyses at each age, provided as a separate excel file. The table lists the eGene names, symbol, and treeQTL p-value, the number of significant eQTLs for the eGene at age 70 (80), and information about the gene validating at age 80 (70). Related to Figure 3.

Table S9: **Age-specific alternative splicing across the transcriptome.** Cluster-and gene-level summary statistics for differential splicing by age analysis in the PIVUS study, provided as a separate excel file. Table lists the cluster name, chromosome, start and end base pair position, the associated gene, and the log likelihood ratio test (LRT) statistic, LRT p-value, and LRT q-value for the age effect. Related to Figure 5.

Table S10: **Gene-ontology enrichment analysis for genes differentially spliced with age.** The table, provided as a separate excel file, lists summary statistics for enriched GO terms, i.e. enriched gene set names, description, size, observed and expected overlap with differentially spliced genes, and enrichment score, p-value, and q-value. Related to Figure 5.

Table S11: **Age-specific genetic regulation of alternative splicing across the transcriptome.** Summary statistics for the sQTL analyses at each age, provided as a separate excel file. The table lists the cluster name, chromosome, start and end base pair position, the associated gene, the treeQTL p-value, the number of significant sQTLs for the intron at age 70 (80), and information about the sQTL validating at age 80 (70). Related to Figure 5.

## Supplemental Figures

**Figure S1:**
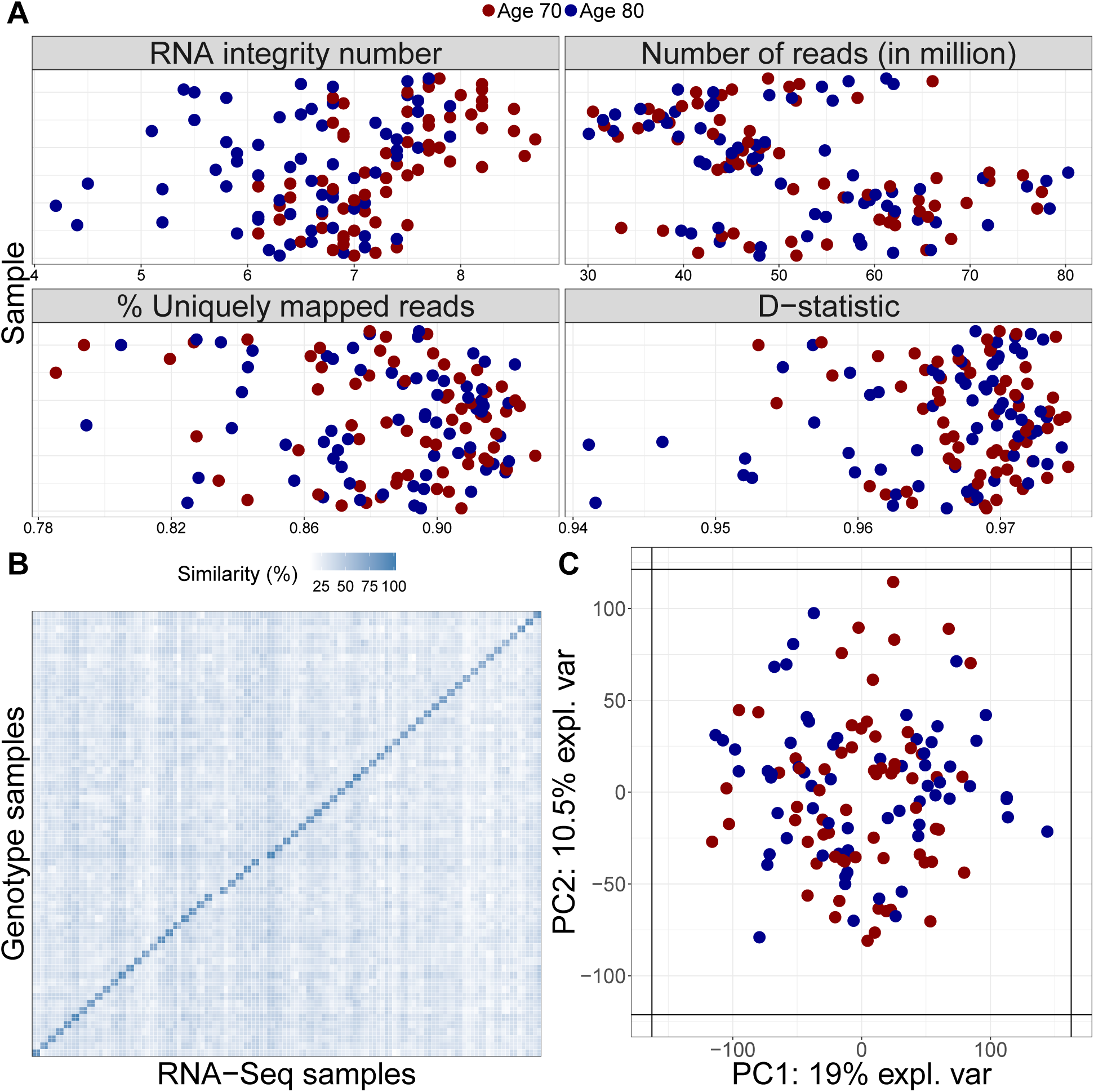
RNA-Seq data quality control. **(A)** Distribution of RNA integrity number, number of sequenced reads (in million), percent of uniquely mapped reads, and D-statistic across samples. The median RNA integrity number across samples was 6.9. All samples had at least 30M reads, at least 80% of their reads mapped uniquely, and their median Spearman expression correlation (D-statistic) with other samples was at least 0.9. **(B)** Concordance between SNP array and RNA-Seq called heterozygous loci. All RNA-seq samples are most similar to their own genotype sample (darker numbers in the diagonal). The four light colors in the diagonals refer to the two individuals (four samples) for which SNP genotypes were not available; these individuals are not similar to the genotype of any other sample. **(C)** Principal component analysis on expression data. No outliers are present based on the two first principal components (PC). Related to Figure 1.

**Figure S2:**
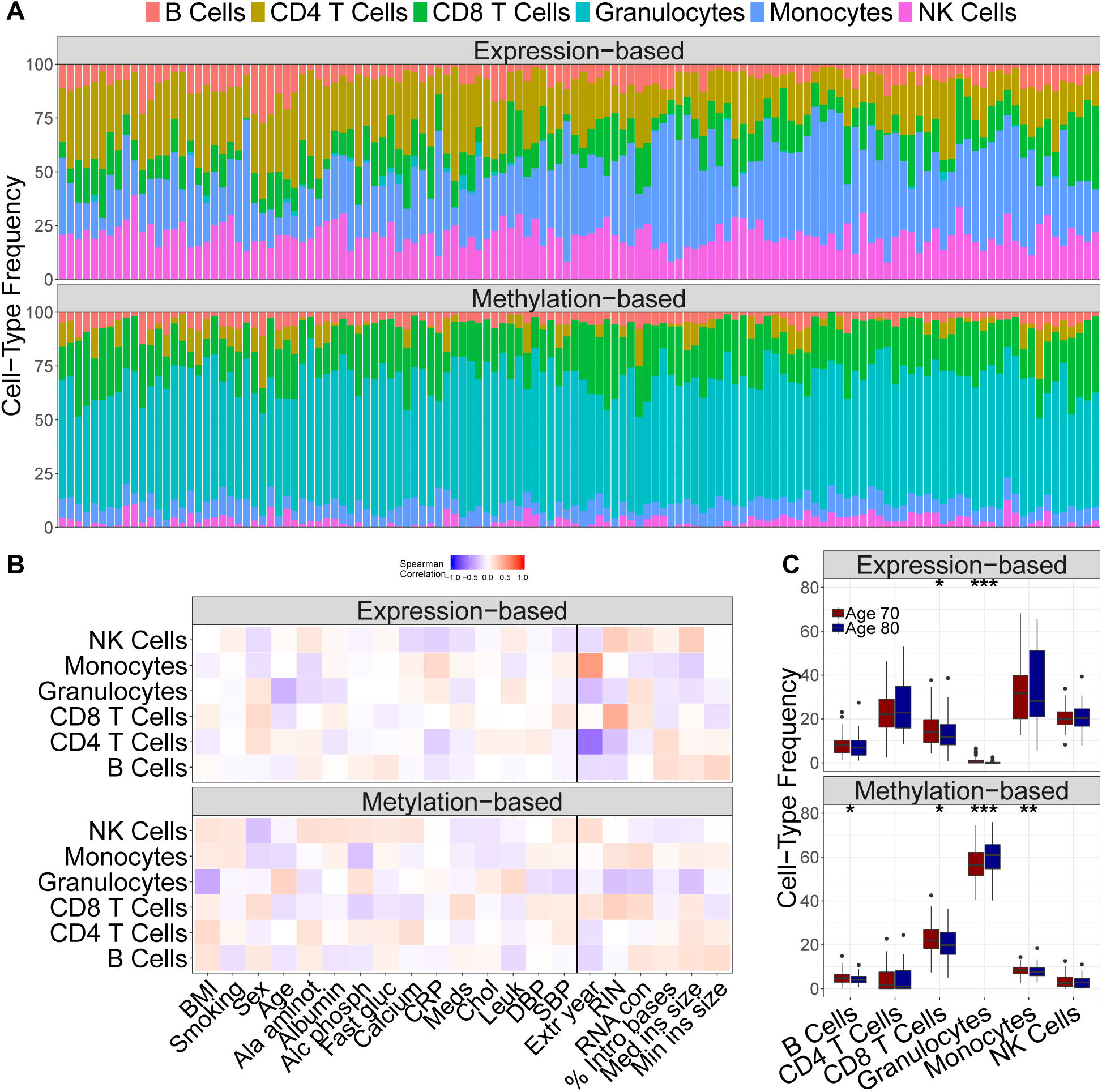
Cell-type deconvolution in whole blood. **(A)** Cell-type frequency estimates (in %). Each column contains relative estimates for each sample. Pairs of consecutive columns are 70-and 80-years-old samples of the same individual. Expression-based estimates are not close to expected values for adults of similar age. **(B)** Correlation of cell-type frequencies with biological and technical covariates (separated by black line). Expression-based estimates are highly correlated with technical covariates while methylation-based estimates are correlated with biological covariates. **(C)** Distribution of estimated cell-type frequencies by age. For expression-based estimates, we see a significant difference with ages for CD4 and CD8 T cells and granulocytes (2-sample Wilcoxon test; *p values* = 0.053,.029, and 6.73 × 10^−5^). For methylation-based estimates, B and CD8 T cells, granulocytes, and monocytes showed a significant difference between the two ages (p-values = 0.018, 0.019, 1.21 × 10^−4^, and 9.29 × 10^−3^). Significance codes: ′ * **′ ≤ 0.001, ′* *I ≤ 0.01, ′*′ ≤ 0.05. Related to Figure 1.

**Figure S3:**
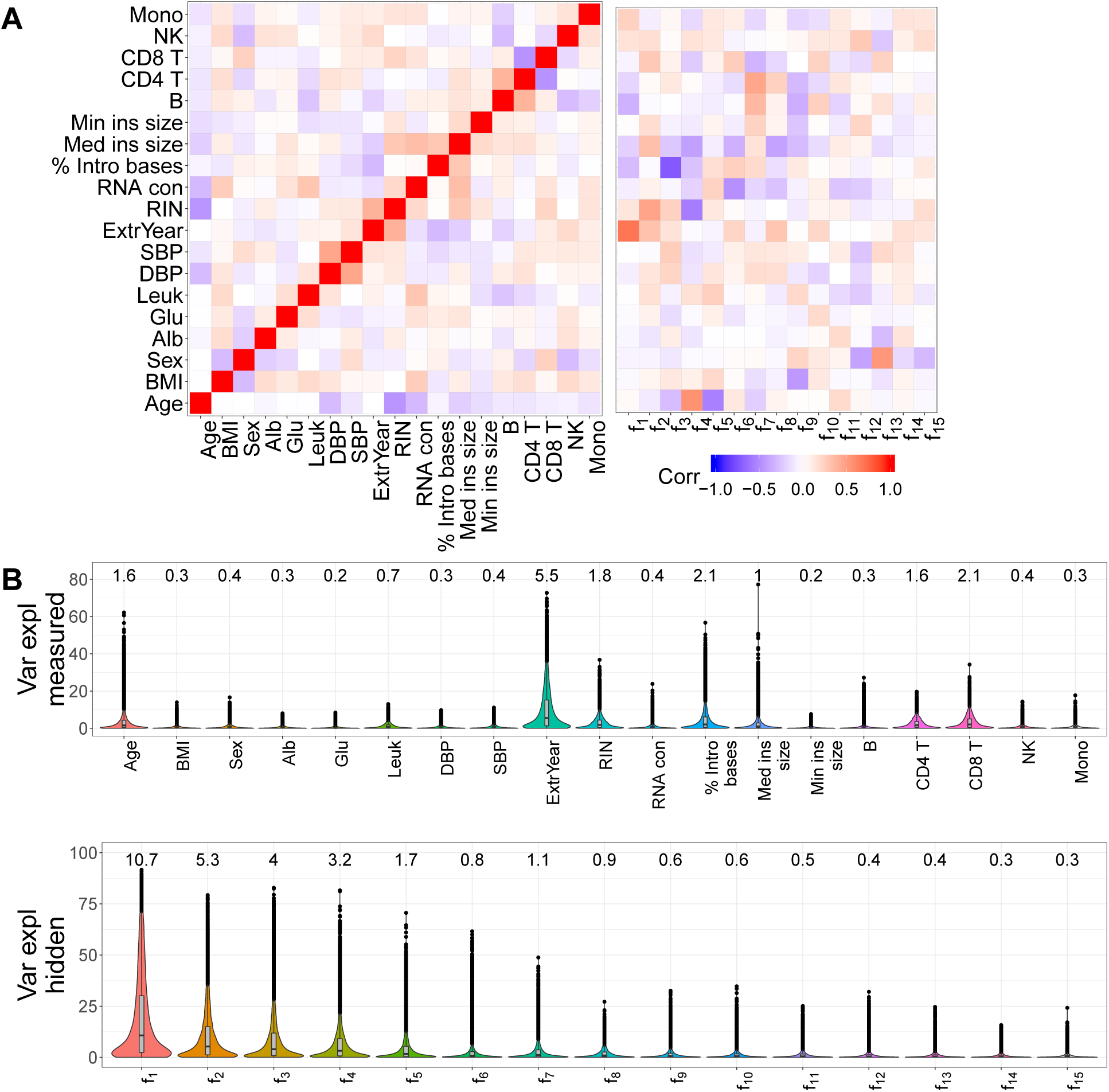
Known and inferred determinants of gene expression variability. **(A)** Correlation between measured covariates and with hidden factors. Age is moderately correlated with RIN (Spearman’s *ρ* =-0.46, p-value = 5.16×10^−8^) and RNA concentration (Spearman’s *ρ* = −0.30, p-value = 4.21×10^−4^). Hidden factors are correlated with several measured factors, e.g. Spearman’s *ρ* = .69 between extraction year and first hidden factor. The fourth and fifth hidden factors are moderately correlated with age (Spearman’s *ρf*_4,*age*_ = 0.56 and *ρf*_5,*age*_ = −0.55). The fourth hidden factor is also correlated with RIN (Spearman’s *ρ* = 0.56), consistent with the fact that age and RIN are moderately correlated. **(B)** Proportion of gene expression variance explained (VE) by measured and hidden factors. VE for each measured factor is estimated by fitting a multiple linear mixed model with all measured factors and age for each gene. Consistent with results in (A), extraction year accounts for the largest proportion of gene expression variance (VE mean= 9.89%, median = 5.54%). Measured covariates are listed in the same order as Table S1. Moreover, the first hidden factor explains about 10% of gene expression variance (median across genes) while the 15th one explains about.3%. Related to Figure 1.

**Figure S4:**
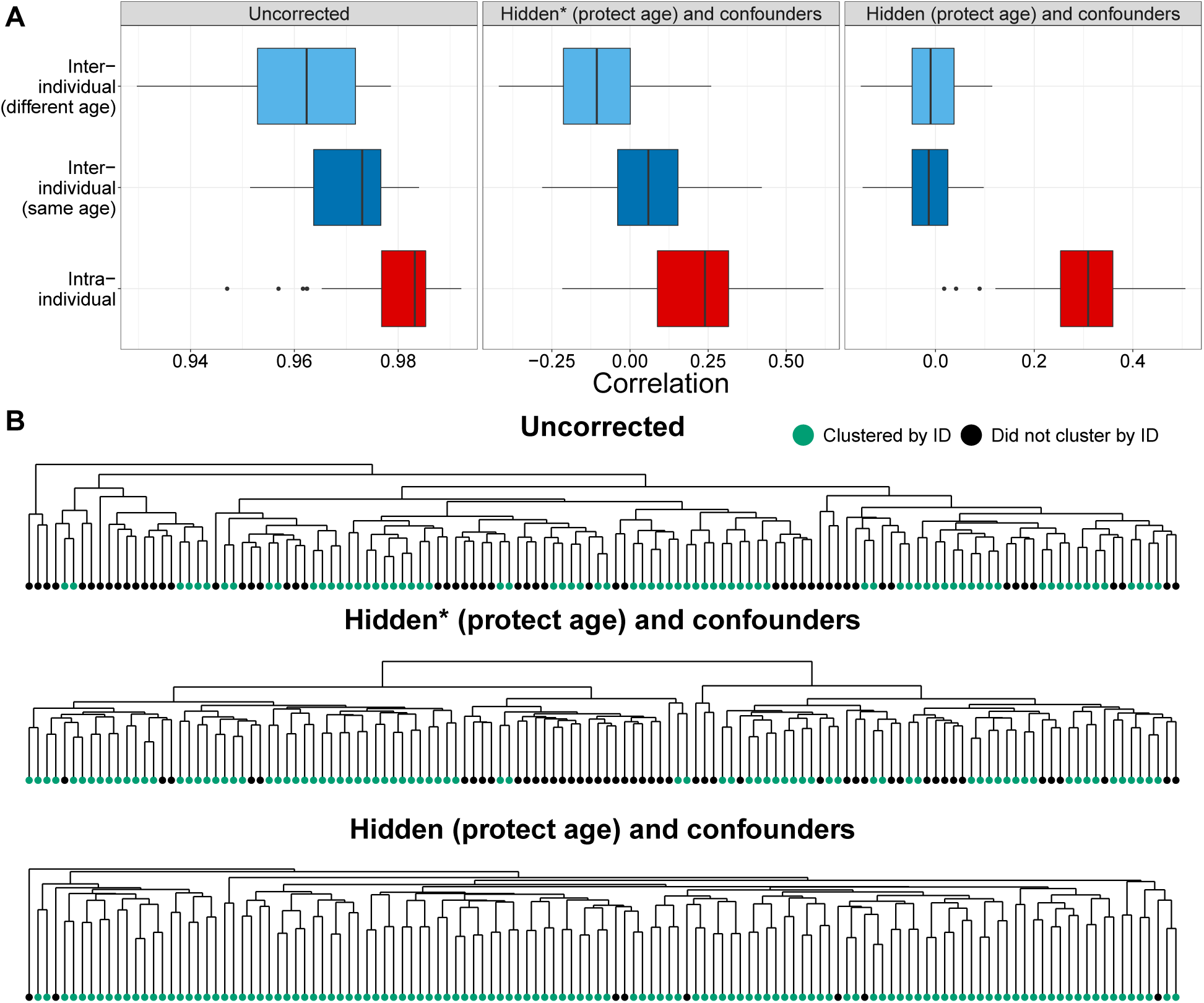
Gene expression within individuals is highly correlated. **(A)** Intra-and inter-individual gene expression correlations (Spearman’s *ρ*; across all genes) based on uncorrected data (Uncorrected), data corrected for confounders and all inferred components of gene expression variability (Hidden (protect age) and confounders) or confounder and only inferred components that are uncorrelated with age (Hidden* (protect age) and confounders). Intra-individual (red) refers to samples of the same individual at the two ages. Inter-individual refers to samples of two different individuals that share age (dark blue) or do not share age (light blue), for 65 randomly sampled pairs of individuals. Even after correcting for global determinants of gene expression variability, the intra-individual correlations are higher than the inter-individual correlations, due to cis genetic and environmental effects that are unique to the individuals. **(B)** Dendrogram of expression-based sample-to-sample distance. Measurements of the same individual at the two ages cluster together (green) for the majority of samples. Labels of nodes denote sample ids; odd number ids are age 70 samples while even are age 80. Pairs of consecutive ids refer to the same individual, e.g. 3 and 4 refer to the same individual at age 70 and 80. Related to Figure 1.

**Figure S5:**
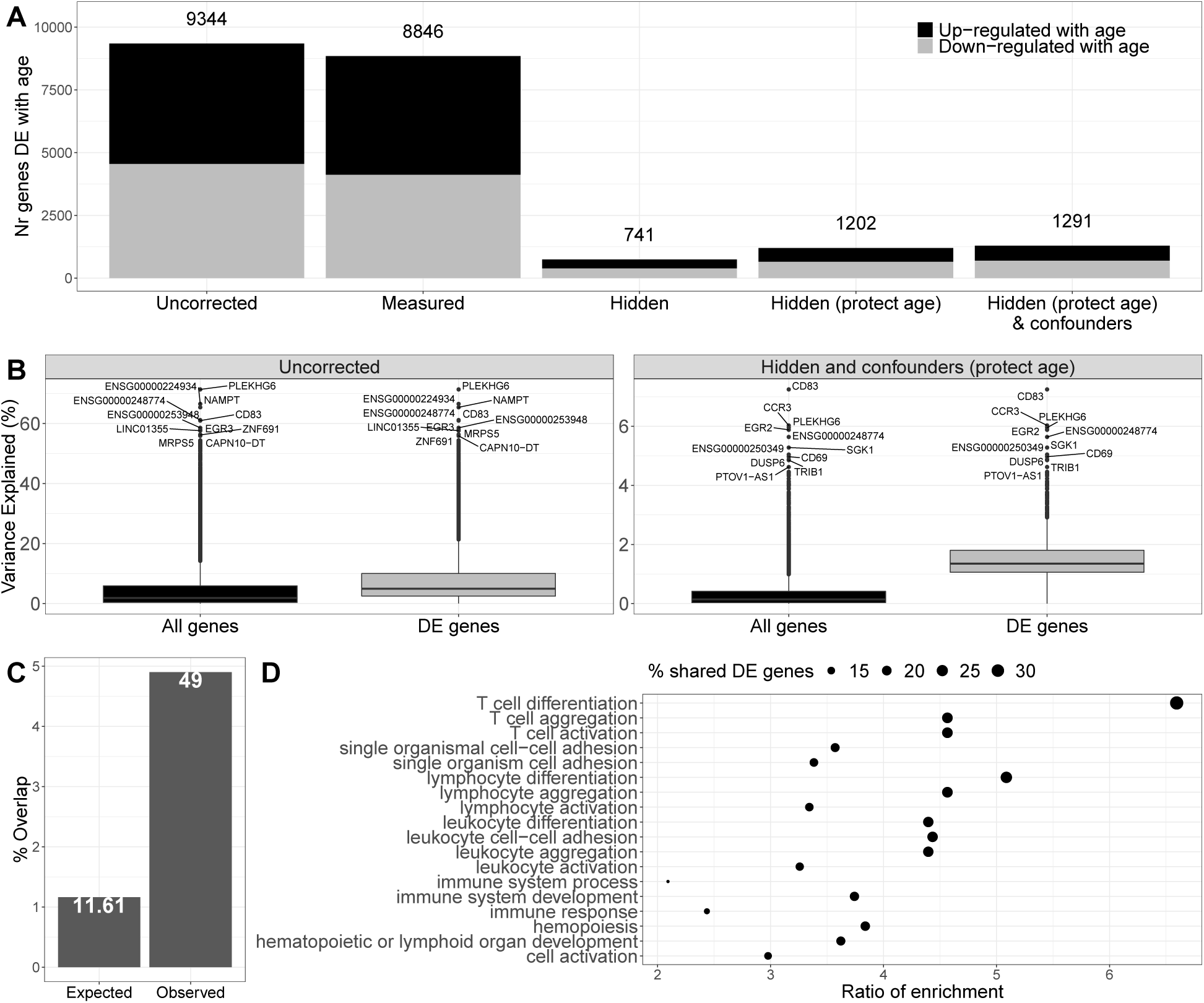
Population-level age-specific expression across the transcriptome. **(A)** Proportion of tested genes differentially expressed (DE) with age and DE genes that show down-regulation with age (FDR 5≤%) for different background noise correction methods. Measured factors are listed in Table S1. Confounders are RIN and RNA-concentration. Numbers on top of bars show number of DE genes. **(B)** Box-plot of expression variance explained by age (in %) across all genes (black) or significantly DE genes (grey). In uncorrected data, age explained, on average, 4.8% of the expression variance of all genes (median=1.84%) and 7.9% (median=4.97%) for genes DE with age. In the data corrected for hidden factors and confounders, age explained a smaller proportion of expression variance (note the difference in y-axis scale), since we removed part of expression variance attributed to age that could be due to confounders. Globally, age explained, on average,.31% (median= .14%) of expression variance, while for genes DE with age, it explained 1.5% (median=1.3%) of expression variability. **(C)** Proportion of overlap between the top 1,000 DE genes in PIVUS with the top 1,000 DE genes from CHARGE and SardiNIA. We found a statistically significant overlap between PIVUS and the other two studies (Fold enrichment = 4.17; Hyper-geometric exact test; *p value* = 1.3 ×10^−16^). **(D)** Gene ontology enrichment analysis for the 49 genes that are in the top 1000 most DE genes across all three studies. Related to Figure 1.

**Figure S6:**
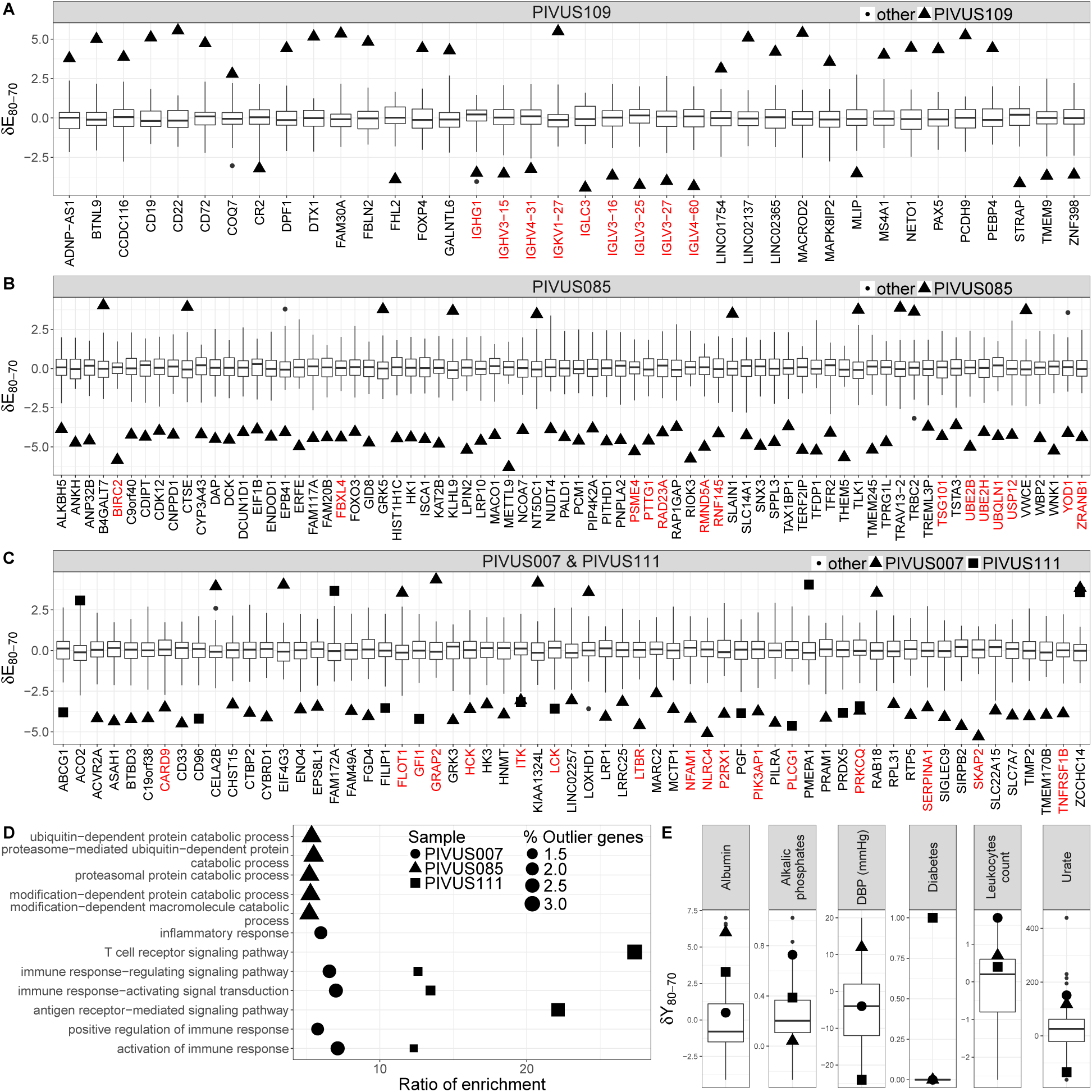
Examples of age-trajectory outliers. **(A-C)** Outlier genes for four different individuals. Y-axis shows the expression differences between two ages (*δE*_80−70_). **(D)** Enrichment for GO biological processes for each individual’s set of outlier genes. **(E)** Outlier phenotypes for the four individuals. Y-axis shows the phenotype differences between two ages (*δY*_80−70_). The larger symbols indicate where the outlier individual is located in the distribution of each gene/phenotype. Related to Figure 2.

**Figure S7:**
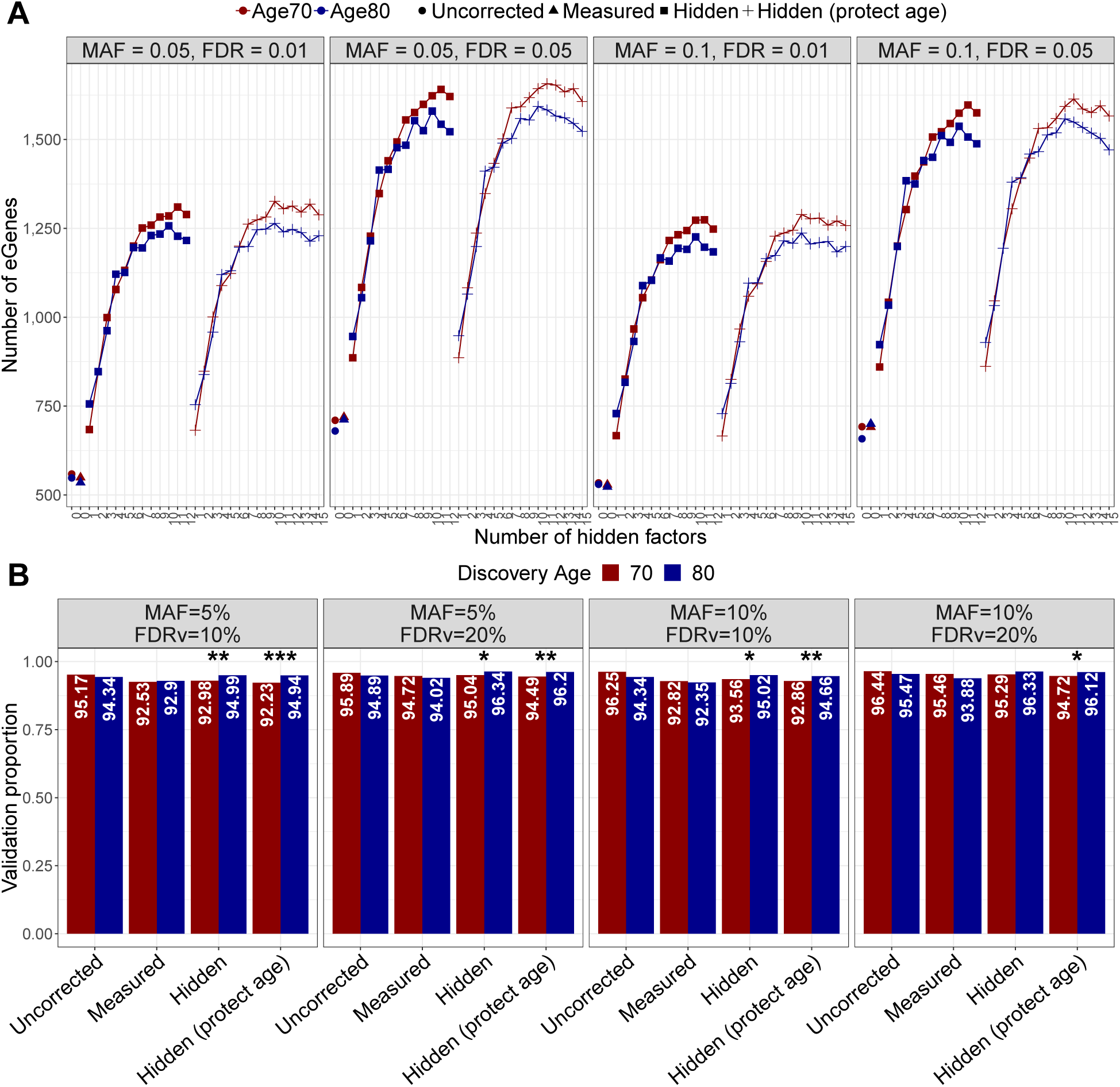
Age-specific genetic regulation across the genome. **(A)** Number of genes with at least one significant eQTL (eGenes) at age 70 (red) and 80 (blue) for different background noise correction strategies (shape), minor allele frequencies (MAF), and FDR thresholds used to identify eGenes and eQTLs**^?^**. We find the largest number of discoveries when we correct for hidden factors. For all MAFs and FDR thresholds we make more discoveries at age 70, compared to age 80. **(B)** Proportion of eGenes discovered at age 70 (80) that validated at age 80 (70) for different background noise correction strategies, validation FDR levels (FDRv), and MAF filters for candidate eQTL at each age. The discovery FDR was 1%. *, **, and *** indicate that the validation proportion at age 70 is lower than the one at age 80 at ≤10%, 5%, and 1% nominal significance levels, respectively. Related to Figure 3.

**Figure S8:**
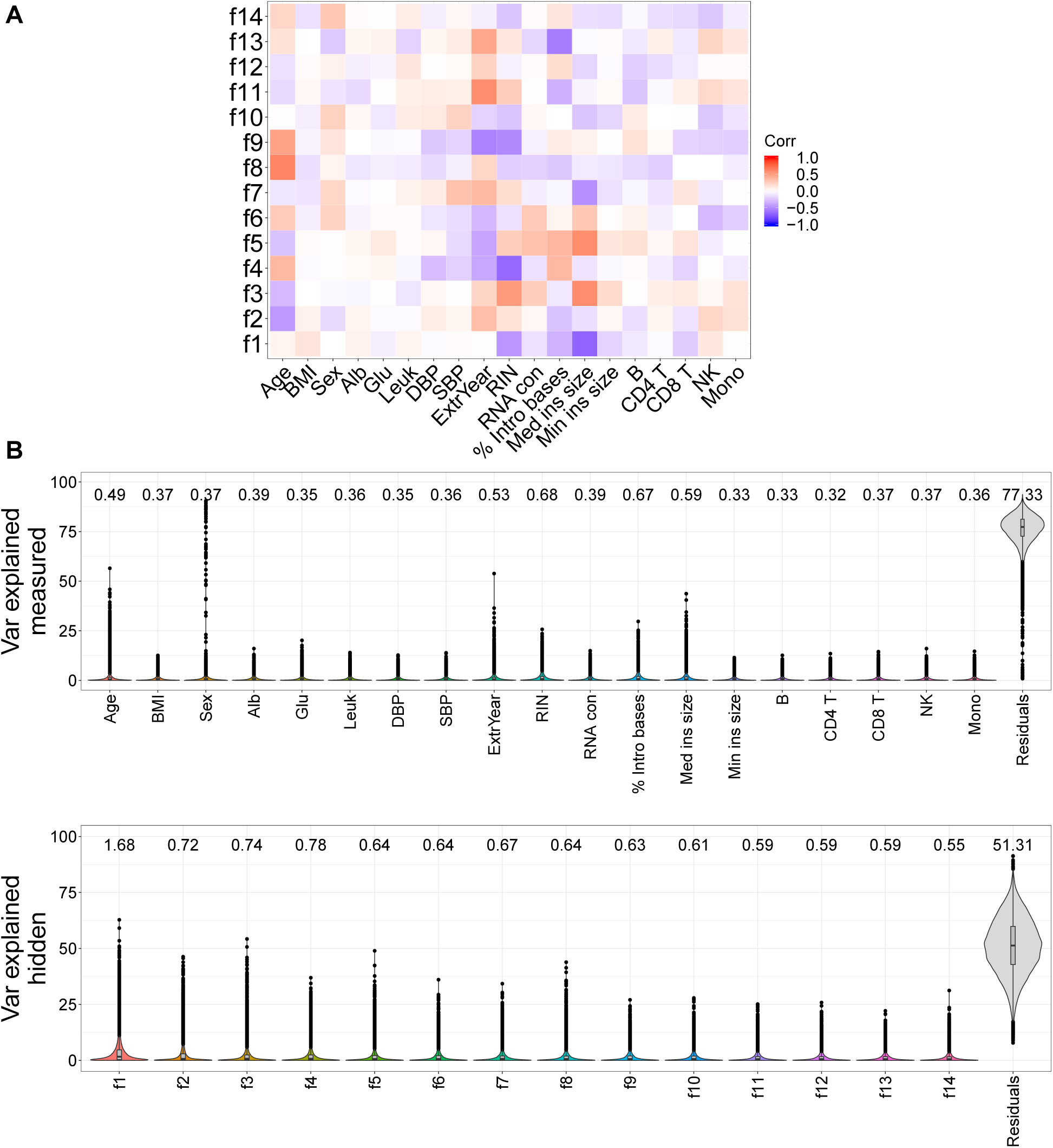
Known and inferred determinants of alternative splicing variability. **(A)** Correlation between measured covariates and alternative splicing hidden factors. Hidden factors are correlated with several measured factors. Several hidden factors are correlated with age, e.g. Spearman’s *ρ*_*f*_8*,age* = 0.53. **(B)** Proportion of alternative splicing variance explained (VE) by measured and hidden factors. VE for each measured or hidden factor is estimated by fitting a multiple linear mixed model with all measured or hidden factors for each gene. VE by the measured or hidden factors for alternative splicing is lower than the VE for expression. This is due to intron excision ratios being internally normalized and thus less affected by technical variability. Related to Figure 5.

